# Kin17 drives dissociation of Mira from the centrosome in neuroblasts by regulating splicing of Flfl

**DOI:** 10.1101/2021.11.03.467193

**Authors:** Marisa Connell, Yonggong Xie, Rui Chen, Sijun Zhu

**Affiliations:** Department of Neuroscience and Physiology, State University of New York Upstate Medical University, Syracuse, NY 13210, USA; GeneScript USA, Inc, Piscataway, NJ 08854, USA

## Abstract

During asymmetric division of *Drosophila* neuroblasts, the fate determinant Prospero and its adaptor Miranda are segregated to the basal cortex through aPKC phosphorylation of Miranda and displacement from the apical cortex. Here we identify Kin17 as a novel regulator of Miranda localization during asymmetric cell division and loss of Kin17 or Protein Phosphatase 4 leads to aberrant localization of Miranda to the centrosome and cytoplasm and Prospero to the centrosome and nucleus. We report that dephosphorylation of Mira by Protein Phosphatase 4 at Serine-96 at the centrosome is required for the proper basal localization of Mira after being phosphorylated at the apical cortex. We further demonstrate that Kin17 regulates Miranda localization by promoting splicing of the transcript of a PP4 component Falafel. Taken together, our work reveals a novel mechanism that ensures proper basal localization of Miranda by preventing its aberrant localization to the centrosome during the asymmetric division.

**Summary:** Connell et al. show that proper segregation of cell fate determinant Miranda requires PP4-mediated dissociation of Miranda from the centrosome/spindle in *Drosophila* neuroblasts, a critical step that has been overlooked previously. Further, they identify a novel regulator of Miranda localization, Kin17, which promotes splicing of the transcript of the PP4 component Falafel.

## Introduction

The generation of diverse cell types from a single population of stem cells is essential for proper development. In the brain, neural stem cells (NSCs) divide to produce neurons and glia. Asymmetric cell division allows for the generation of cells of different fates from one parental cell by producing daughters of different protein content, fate, niche, and/or size. This process must be tightly regulated to ensure the proper balance of stem and differentiating cells, and defects in this process can lead to overproliferation or premature differentiation of the stem cell population, both of which can have detrimental effects on the organism, such as cancer or developmental disorders (Gómez-López *et al*, 2014).

*Drosophila* NSCs, termed neuroblasts (NBs), have provided a useful model for the study of the mechanisms that regulate asymmetric cell division as many of the key components are conserved in mammalian systems including humans (Homem & Knoblich, 2012). In developing *Drosophila* brains, most NBs (type I NBs) divide asymmetrically to generate a self-renewing neuroblast and a differentiating ganglion mother cell (GMC) (Homem & Knoblich, 2012). GMCs divide a single time to produce neurons and glia. The identity of the GMC is established by the specific segregation of the fate determinants, Prospero (Pros) (Hirata *et al*, 1995; Knoblich *et al*, 1995), Numb (Knoblich *et al*, 1995), and Brain tumor (Brat)(Bello *et al*, 2006; Betschinger *et al*, 2006; Lee *et al*, 2006) to the GMC during mitosis. In order to deliver these factors to the GMC, neuroblasts establish a cell polarity axis during mitosis through the recruitment of the Par protein complex to the apical pole of the cell (Betschinger *et al*, 2003; Knoblich, 2010). This apical complex establishes cell polarity by excluding the localization of the fate determinants from the apical pole, leading to basal localization of these factors.

One of the primary factors that must be excluded from the apical pole of neuroblasts is the adaptor protein, Miranda (Mira), which localizes *pros* mRNA (Schuldt *et al*, 1998), Pros (Ikeshima-Kataoka *et al*, 1997; Shen *et al*, 1997), Staufen (Matsuzaki *et al*, 1998), and Brat (Betschinger *et al*, 2006) to the basal pole and ensures their proper segregation to the GMC. Prior to the onset of mitosis, Mira localizes to the entire cell cortex through an interaction between its Basic and Hydrophobic (BH) motif and the cell membrane (Bailey & Prehoda, 2015). At the onset of mitosis, Inscuteable and Bazooka localize to the apical pole and recruit the serine/threonine kinase atypical Protein Kinase C (aPKC) through its activator Par6 (Schober *et al*, 1999; Wodarz *et al*, 1999; Petronczki & Knoblich, 2001; Albertson & Doe, 2003). aPKC phosphorylates Mira within the BH motif at Serine-96 (S96), Threonine-194, Serine-195, Threonine-205, and Serine-206, breaking the interaction between Mira and the apical cell cortex, leading to localization to the basal domain (Atwood & Prehoda, 2009; Bailey & Prehoda, 2015). Although aPKC phosphorylates multiple residues, a phosphomimetic mutation of S96, Mira^S96D^ is almost entirely displaced from the cortex when combined with aPKC knockdown, indicating that phosphorylation of this residue is sufficient for displacement from the cortex. This leads to the question of how phosphorylated Mira can localize to the basal cortex during cell division. A previous study reported that depletion of Falafel (Flfl), the targeting subunit of Protein Phosphatase 4 (PP4), leads to a reduction in Mira localization to the basal domain (Sousa-Nunes *et al*, 2009), implying that Mira may need to be dephosphorylated before interacting with the basal domain. However, while Flfl has been shown to physically interact with Mira (Sousa-Nunes *et al*, 2009), PP4 has only been shown to be required for the dephosphorylation of threonine-591, which is required for the cortical localization of Mira prior to its phosphorylation by aPKC (Zhang *et al*, 2016). Therefore, although how Mira is phosphorylated by aPKC and displaced from the apical cortex has been well studied, it remains unclear how Mira localizes to the basal cortex after aPKC phosphorylation.

In addition to its cortical localization, Mira has been shown to localize to the centrosome of wild-type neuroblasts during prophase (Mollinari *et al*, 2002) and to the mitotic spindle in syncytial embryos and various mutants, including Mira^S96D^ (Mollinari *et al*, 2002; Hannaford *et al*, 2018; Albertson & Doe, 2003; Atwood *et al*, 2007; Petritsch *et al*, 2003). These data suggest that there may be a mechanism by which Mira localizes to the mitotic spindle and is removed during mitosis in *Drosophila* neuroblasts to ensure the proper segregation of Mira and its cargo protein Pros to the daughter cells. However, the mechanism that regulates the localization of Mira to the centrosome/mitotic spindle has not been investigated.

Kin17 is a DNA and RNA binding protein that is essential for DNA replication and DNA damage repair (Angulo *et al*, 1991; Miccoli *et al*, 2005; Masson *et al*, 2003; Pinon-Lataillade, 2004). It has also potentially been identified as interacting with the spliceosome through mass spectrometry in both human and *Drosophila* cells, though the interaction may be transcript specific (Herold *et al*, 2009; Gaspar *et al*, 2021; Rappsilber *et al*, 2002). We have identified Kin17 as a novel factor required for the localization of Mira during asymmetric cell division. Kin17 is required for splicing of the *flfl pre-mRNA* and its loss leads to reduced Flfl levels and defects in Mira localization, including an accumulation of Mira at the centrosome. We found that PP4 at the centrosome directly dephosphorylates Mira at S96 to ensure dissociation of Mira from the centrosome/spindle and segregation of Mira and Pros to the GMC.

## Results

### Kin17 knockdown leads to reduced brain size and a reduced mitotic rate in neuroblasts

We identified Kin17 in a screen for factors that affect *Drosophila* neuroblast development. Kin17 knockdown using the pan-neuroblast driver, *insc-Gal4*, led to larval lethality and reduced brain size compared to control larvae (Fig. 1A). This decrease in brain size was not due to a reduction in neuroblast numbers, as we observed no significant difference in the number of central brain neuroblasts (Figs. 1B,C). However, we observed a significant decrease in the mitotic rate of neuroblasts in *Kin17 RNAi* brains (Figs. 1D,E) and a corresponding reduction in GMCs (Pros+, Elav-) and newly born neurons (Pros+, Elav+) (Fig. 1F). These phenotypes were rescued by expressing wild-type Kin17, ruling out off-target effects of the RNAi (Figs. 1A, D-F). These results indicate Kin17 is required for normal brain development and the promotion of mitosis in neuroblasts.

**Figure 1.**
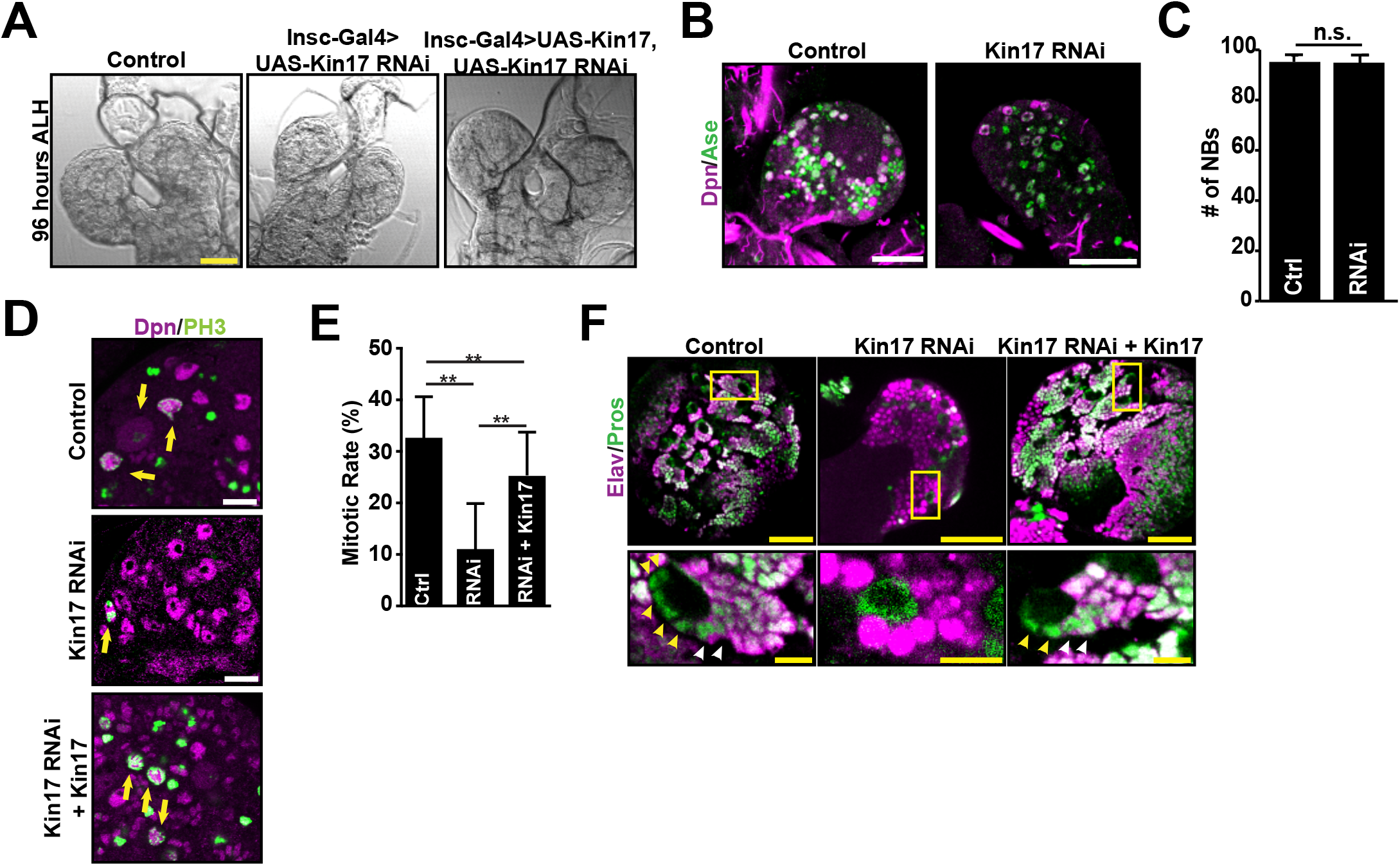
Kin17 knockdown leads to a reduction in brain size and defects in asymmetric cell division. (**A**) *Kin17 RNAi* brains are smaller at 96 hours after larval hatching (ALH) than wild-type brains. Scale bar, 50 μm. (**B**) Kin17 knockdown does not alter the number of neuroblasts in the larval brain. Scale bars, 50 μm. (**C**) Quantification of neuroblast number in control (n=11) and *Kin17 RNAi* (n=10) brains. Error bars represent one standard deviation (1 SD); n.s., not significant. (**D**) PH3 staining in control, *Kin17 RNAi*, and Kin17 rescue neuroblasts. Arrows indicate mitotic neuroblasts with positive PH3 staining. Scale bars, 20 μm. (**E**) Quantification of mitotic rate in control (n=19), *Kin17 RNAi* (n=14), and rescue (n=19) neuroblasts. Error bars, 1 SD; **, *p* < 0.01. (**F**) Kin17 knockdown leads to a reduction of Pros+ progeny. Yellow arrowheads indicate Pros+ GMCs and white arrowheads indicate Pros+, Elav+ neurons. Scale bars, 50 μm.

### Accumulation of nuclear Pros leads to Kin17 knockdown phenotypes

It has been shown previously that expression of nuclear Pros promotes cell cycle exit of GMCs (Colonques *et al*, 2011) and accumulation of Pros in the nucleus of neuroblasts leads to quiescence or premature differentiation (Lai & Doe, 2014). Thus we examined if the reduction in the mitotic rate of neuroblasts in Kin17 knockdown was caused by Pros localization to the nucleus. We observed that 38.5± 6.7% of *Kin17 RNAi* neuroblasts had Pros localized to the nucleus (Figs. 2A,B), which was never observed in control neuroblasts, suggesting that the reduced mitotic rate may be due to accumulation of nuclear Pros.

**Figure 2.**
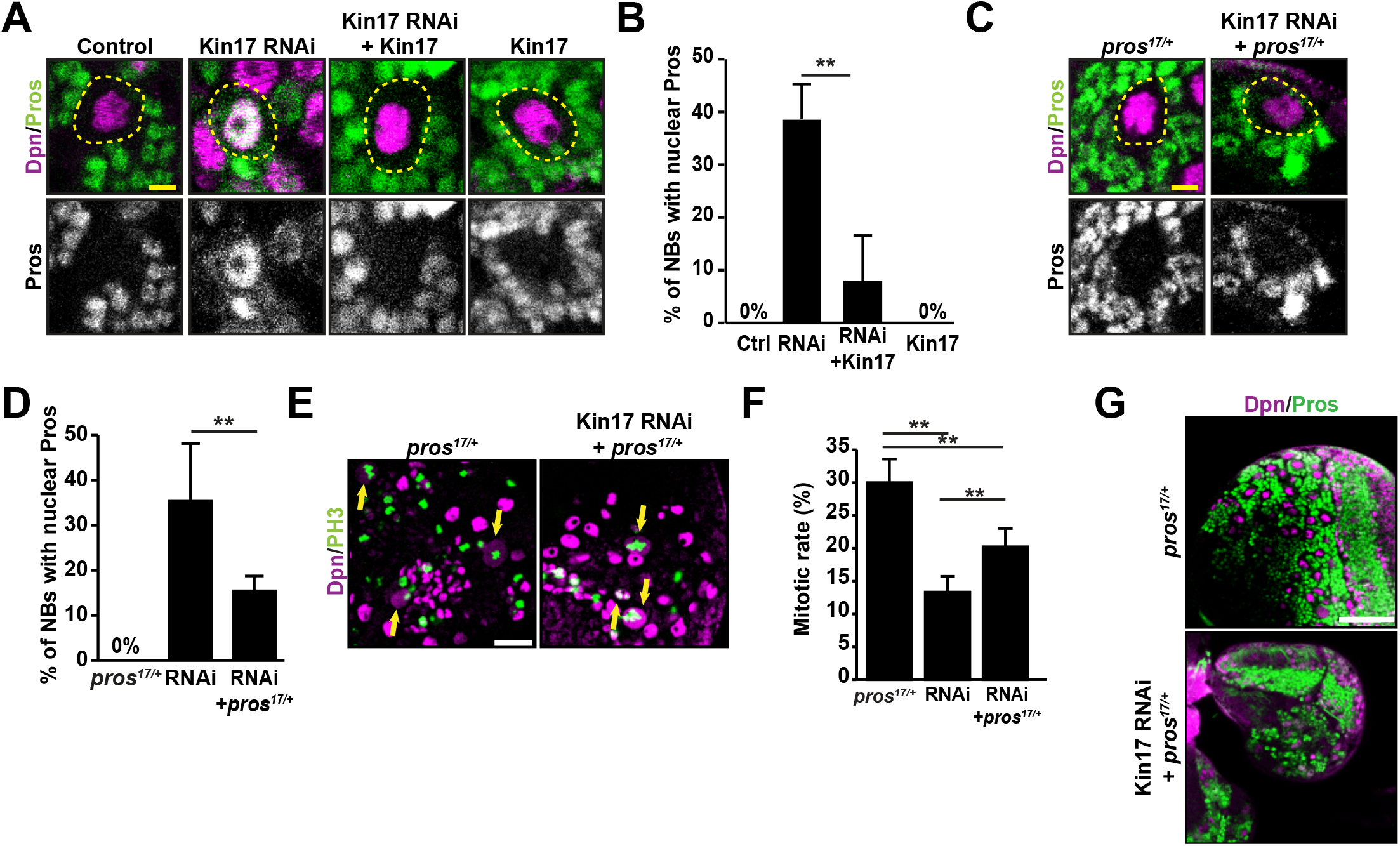
Kin17 knockdown leads to defects in Pros localization. (**A**) Pros staining in control, *Kin17 RNAi*, Kin17 overexpression, and Kin17 rescue brains. Note: Pros is detected in the nucleus of *Kin17 RNAi* neuroblasts. Dotted lines indicate neuroblasts. Scale bars, 5 μm. (**B**) Quantification of Pros localization to the nucleus in control (n=8), *Kin17 RNAi* (n=10), Kin17 rescue (n=10), and Kin17 overexpression (n=11) brains. Error bars, 1 SD; **, *p* < 0.01.(**C**) Pros nuclear localization resulting from Kin7 RNAi is reduced in the *pros*^*17/+*^ background. Dotted lines indicate neuroblasts. Scale bar, 5 μm. (**D**) Quantification of Pros nuclear localization in *pros*^*17/+*^ (n=6), *Kin17 RNAi* (n=10), and *Kin17 RNAi pros*^*17/+*^ (n=8) brains. Error bars, 1 SD; **, p<0.01. (**E**) PH3 staining in *pros*^*17/+*^, *Kin17 RNAi*; *pros*^*17/+*^ neuroblasts. Arrows indicate PH3^+^ mitotic neuroblasts. Scale bar, 20 μm. (**F**) Quantification of mitotic rate in *pros*^*17/+*^ (n=9), *Kin17 RNAi* (n=8), and *Kin17 RNAi; pros*^*17/+*^ (n=11) brains. Error bars, 1 SD; **, *p*<0.01. (**G**) Pros^+^ neurons are partially restored in *Kin17 RNAi; pros*^*17/+*^ brains. Scale bar, 50 μm.

To determine if the aberrant nuclear Pros is responsible for the Kin17 knockdown phenotypes, we examined if reducing Pros levels would alleviate the Kin17 phenotypes. To this end, we knocked down Kin17 in animals heterozygous for the *pros*^*17*^ allele, a null allele that has been reported to lead to a loss of *pros* mRNA (Doe *et al*, 1991). In the *Kin17 RNAi*, *pros*^*17/+*^ background, Pros nuclear localization was significantly reduced (Figs. 2C,D), and as expected, the mitotic rate was partially rescued to 20.2 ± 2.8% compared to 34.3% in control and 13.4% in *Kin17 RNAi* (Figs. 2E,F), and the loss of Pros^+^ neurons was also largely rescued (Fig. 2G). These data suggest that the Kin17 phenotypes are due to mislocalization of Pros to the nucleus and that Kin17 is required for proper localization of Pros in neuroblasts.

### Pros accumulates at the centrosome in Kin17 knockdown neuroblasts

In addition to the nuclear localization of Pros, we observed bright dots of Pros staining in *Kin17 RNAi* and *Kin17 RNAi; pros*^*17/+*^ neuroblasts, which co-localized with pericentrin, indicating this was centrosomal localization of Pros (Figs.3A-C). In control neuroblasts, centrosomal Pros appeared to be cell cycle-dependent, with a peak occurring in prophase which had 51.7% of neuroblasts exhibiting centrosomal Pros, while centrosomal localization of Pros was consistently observed in 50-60% of Kin17 knockdown neuroblasts throughout the cell cycle. Additionally, in control neuroblasts, Pros primarily localized to the apical centrosome, while in *Kin17 RNAi; pros*^*17/+*^, Pros localized to both apical and basal centrosomes in about half the cells (Fig. 3C). However, a basal crescent of Pros was still observed in 100% of Kin17 knockdown neuroblasts at metaphase as in the wild-type control (Fig. 2H. Please also see below, Fig. 3E).

**Figure 3.**
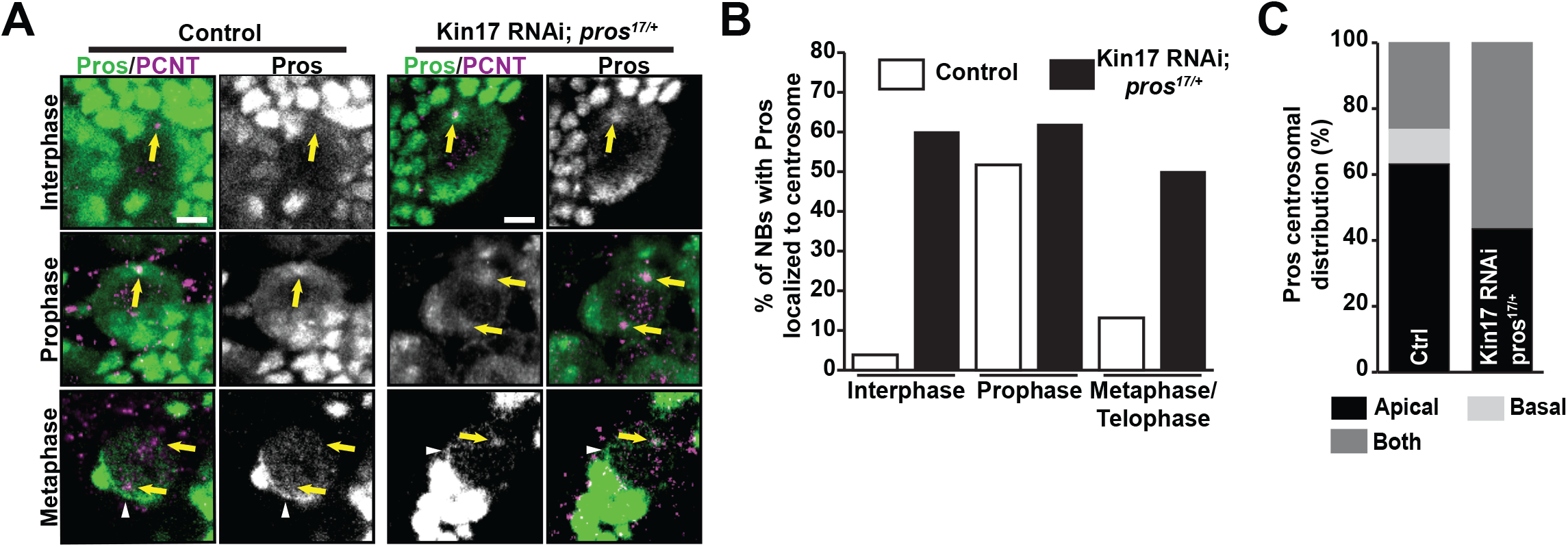
Kin17 knockdown leads to centrosomal localization of Pros. (**A**) Pros localization to the centrosome (marked by Pericentrin (PCNT)) in control and *Kin17 RNAi*; *pros*^*17/+*^ neuroblasts. Arrows indicate centrosome and arrowheads indicated basal crescent. Scale bars, 5 μm. (**B**) Quantification of Pros localization to the centrosome in control and *Kin17 RNAi*; *pros*^*17/+*^ neuroblasts throughout the cell cycle. Interphase, n=26 NBs (ctrl), 25 (*Kin17 RNAi*; *pros*^*17/+*^); Prophase, n=29 (ctrl), 22 (*Kin17 RNAi; pros*^*17/+*^); Metaphase/Telophase; n=24 (ctrl), 16 (*Kin17 RNAi*; *pros*^*17/+*^). (**C**) Quantification of Pros localization to the apical and/or basal centrosome in control (n=19) and *Kin17 RNAi*; *pros*^*17/+*^ (n=23) neuroblasts.

### Kin17 is required for the proper localization of Mira throughout the cell cycle

As Mira, the adaptor protein of Pros, has been shown to localize to the centrosome in prophase (Mollinari *et al*, 2002), we asked if there were changes in the localization of Mira. In wild-type neuroblasts, Mira localized to the cortex as previously described: uniformly cortical in interphase and polarized at the basal pole beginning in prophase (Fig 4A, S1A). In addition, we observed that Mira localized to the centrosome, as indicated by colocalization with γ-tubulin, in 56.9% and 29.8% of neuroblasts at prophase and prometaphase/metaphase, respectively (Figs. 4A,D), which is consistent with a previous report (Mollinari *et al*, 2002). Since Kin17 knockdown neuroblasts have a low mitotic rate, we quantified Mira localization at different stages of the cell cycle in *Kin17 RNAi; pros*^*17/+*^ neuroblasts, as Pros is not required for proper Mira localization (Shen et al., 1997). In *Kin17 RNAi; pros*^*17/+*^ neuroblasts, we found that localization of Mira to the cortex in neuroblasts was reduced and 24.8% and 9.1% of *Kin17 RNAi; pros*^*17/+*^ neuroblasts had completely cytoplasmic localization of Mira at interphase and prometaphase/metaphase, respectively, as well as reduced basal cortical localization in 20-30% of prophase and metaphase neuroblasts compared to control neuroblasts. (Figs. S1A,B). More strikingly, we observed that Mira was consistently localized to the centrosome and/or spindle microtubules in 50-70% of *Kin17 RNAi; pros*^*17/+*^ neuroblasts at all stages of the cell cycle (Figs. 4C,D), which is in stark contrast to wild type neuroblasts. Furthermore, we found 40-45% of Kin17 knockdown neuroblasts had Mira localized to both basal and apical centrosomes, while Mira was almost completely localized to the apical centrosome in control neuroblasts (Fig. S1C). Besides the aberrant centrosomal localization, about 25% of Kin17 knockdown neuroblasts also exhibited Mira localization to the mitotic spindle at anaphase/telophase, which was only observed in 3-6% of neuroblasts in the wild-type control or *pros*^*17/+*^ heterozygous brains. We confirmed that the aberrant centrosomal localization of Mira was not due to the presence of the *pros*^*17*^ allele, as Mira localized to the centrosome in 56.9% of interphase *Kin17 RNAi* neuroblasts (Figs. S1D,E).

**Figure 4.**
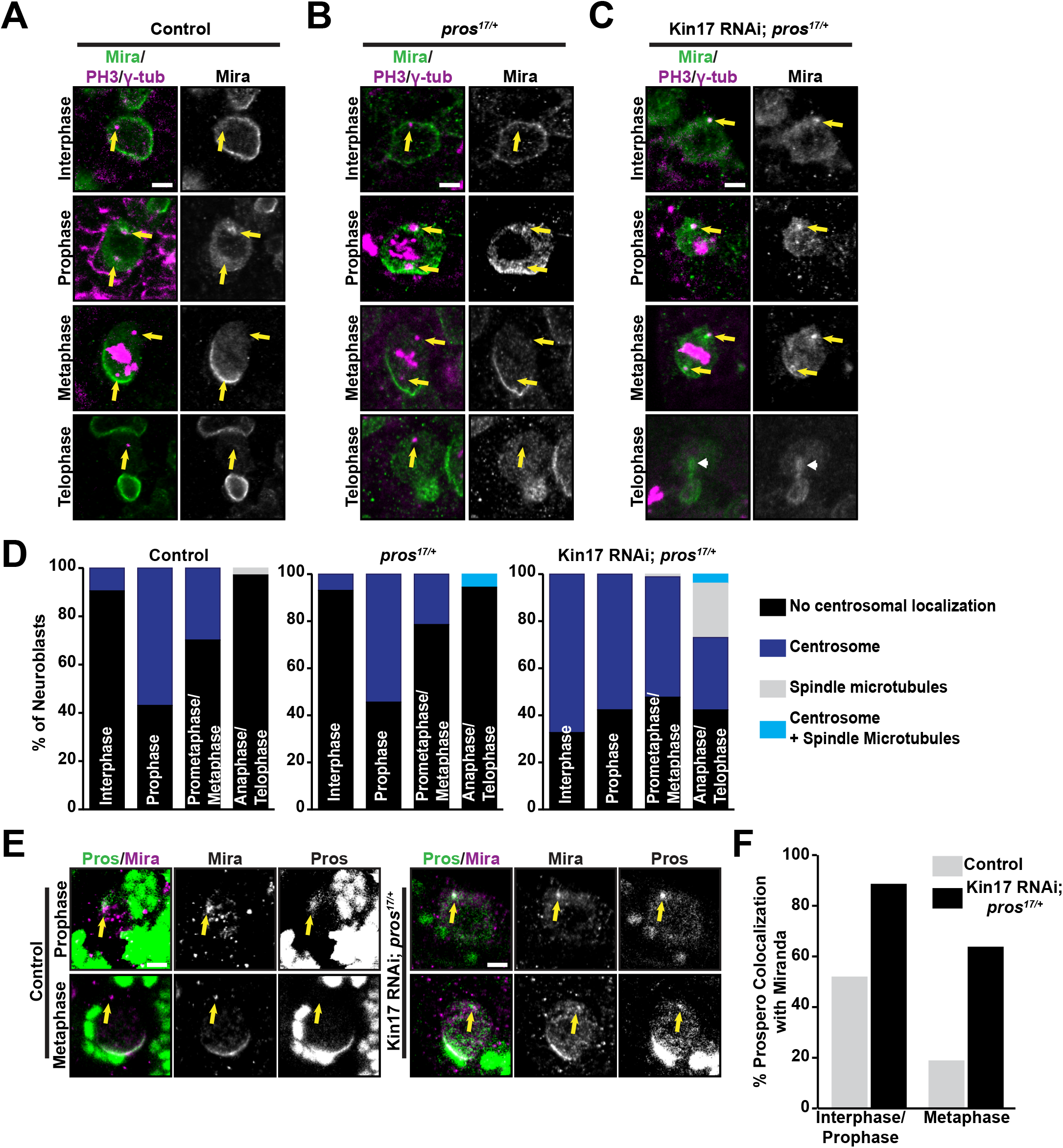
Kin17 knockdown leads to mislocalization of the Pros adaptor Mira. (**A**) Localization of Mira in control neuroblasts throughout the cell cycle. Centrosomal localization of Mira at the prophase is indicated by arrows. Scale bar, 5 μm. (**B**) Localization of Mira in *pros*^*17/+*^ neuroblasts throughout the cell cycle. Scale bar, 5 μm. (**C**) Localization of Mira in *Kin17 RNAi; pros*^*17/+*^ neuroblasts throughout the cell cycle. Note that centrosomal localization of Mira can be observed at all different stages of the cell cycle (arrows). Arrow indicates spindle microtubules. Scale bar, 5 μm. (**D**) Quantification of Mira localization to the centrosome/spindle during the cell cycle in control, *pros*^*17/+*^, and *Kin17 RNAi; pros*^*17/+*^ neuroblasts. Interphase, n=74 NBs (ctrl), 101 (*pros*^*17/+*^), 101 (*Kin17 RNAi; pros*^*17/+*^); Prophase, n=63 (ctrl), 57 (*pros*^*17/+*^), 85 (*Kin17 RNAi; pros*^*17/+*^); Prometaphase/Metaphase, n=57 (ctrl), 56 (*pros*^*17/+*^), 98 (*Kin17 RNAi; pros*^*17/+*^); Anaphase/Telophase, n=35 (ctrl), 18 (*pros*^*17/+*^), 26 (*Kin17 RNAi; pros*^*17/+*^). (**E**) Kin17 knockdown leads to colocalization of Pros with Mira at the centrosome during interphase and metaphase. Arrows indicate colocalization of Mira and Pros at the centrosome. Scale bars, 10 μm. (**F**) Quantification of Pros colocalization with Mira at the centrosome. Interphase/Prophase, n=62 NBs (ctrl), 23 (*Kin17 RNAi; pros*^*17/+*^); Metaphase, n=16 (ctrl), 11 (*Kin17 RNAi; pros*^*17/+*^).

To determine if the centrosomal localization of Pros in Kin17 knockdown neuroblasts was due to the mislocalization of Mira on the centrosome, we co-stained for Mira and Pros. We observed that in Kin17 knockdown when Mira localized to the centrosome, Pros was colocalized to the centrosome 88.5% of the time in interphase/prophase and 63.6% of the time in prometaphase/metaphase, and Pros did not localize to the centrosome in the absence of centrosomal Mira (Figs. 4E,F), indicating that the centrosomal localization of Pros and Mira are correlated and that the centrosomal localization of Pros might be dependent on Mira. Defects in Mira localization to the basal cortex could be due to factors generally affecting basal localization of fate determinants. Therefore, we looked at localization of the fate determinant Numb and found it was not altered in Kin17 knockdown (Figs. S2A,B), indicating that. Kin17 specifically regulates Mira localization. Taken together, our data indicate that Kin17 regulates the localization of Mira at the centrosome in a cell cycle-dependent manner and that aberrant Mira localization correlates with localization of Pros to the nucleus.

### Mira phosphorylation state influences the localization of Mira to the centrosome during asymmetric cell division

As localization of Mira to the centrosome in wild-type neuroblasts begins in prophase when aPKC is recruited to phosphorylate Mira, we hypothesized that phosphorylation of Mira by aPKC leads to the localization of Mira to the centrosome. It has been demonstrated that Serine-96 (S96) is phosphorylated by aPKC and this phosphorylation is required and sufficient for the displacement of Mira from the cell cortex (Atwood & Prehoda, 2009; Bailey & Prehoda, 2015; Hannaford *et al*, 2018). To directly assess if the phosphorylation state of S96 is involved in regulating the localization of Mira to the centrosome, we utilized alleles of *mira*, including wild-type (*mira*^*WT*^), phosphodead (*mira*^*S96A*^), and phosphomimetic (*mira*^*S96D*^) alleles (Hannaford *et al*, 2018), which are tagged with HA and mCherry and can therefore be distinguished from endogenous Mira. We observed that Mira^WT^ localizes similarly to endogenous Mira, except that in anaphase/telophase, Mira^WT^ was enriched at the centrosome and/or mitotic spindle in about 20% of neuroblasts (Figs. S3A). For Mira^S96D^, we observed an increase in cytoplasmic localization at interphase and prophase (Fig. S3A), and increased centrosomal localization through the entire cell cycle (Figs. 5A, B). However, unlike Mira^WT^, which localized only to the apical centrosome in 67.7% of neuroblasts, Mira^S96D^ localized to both apical and basal centrosomes in 54.2% of neuroblasts (Fig. S3B). Mira^S96A^ remained localized to the entire cortex through the cell cycle as expected(Figs. 5A, S3A), and when compared to Mira^WT^, localization to the centrosome was reduced at all stages except for anaphase/telophase (Figs. 5A,B). These data suggest that phosphorylation of S96 of Mira is required for localization of Mira to the centrosome.

**Figure 5.**
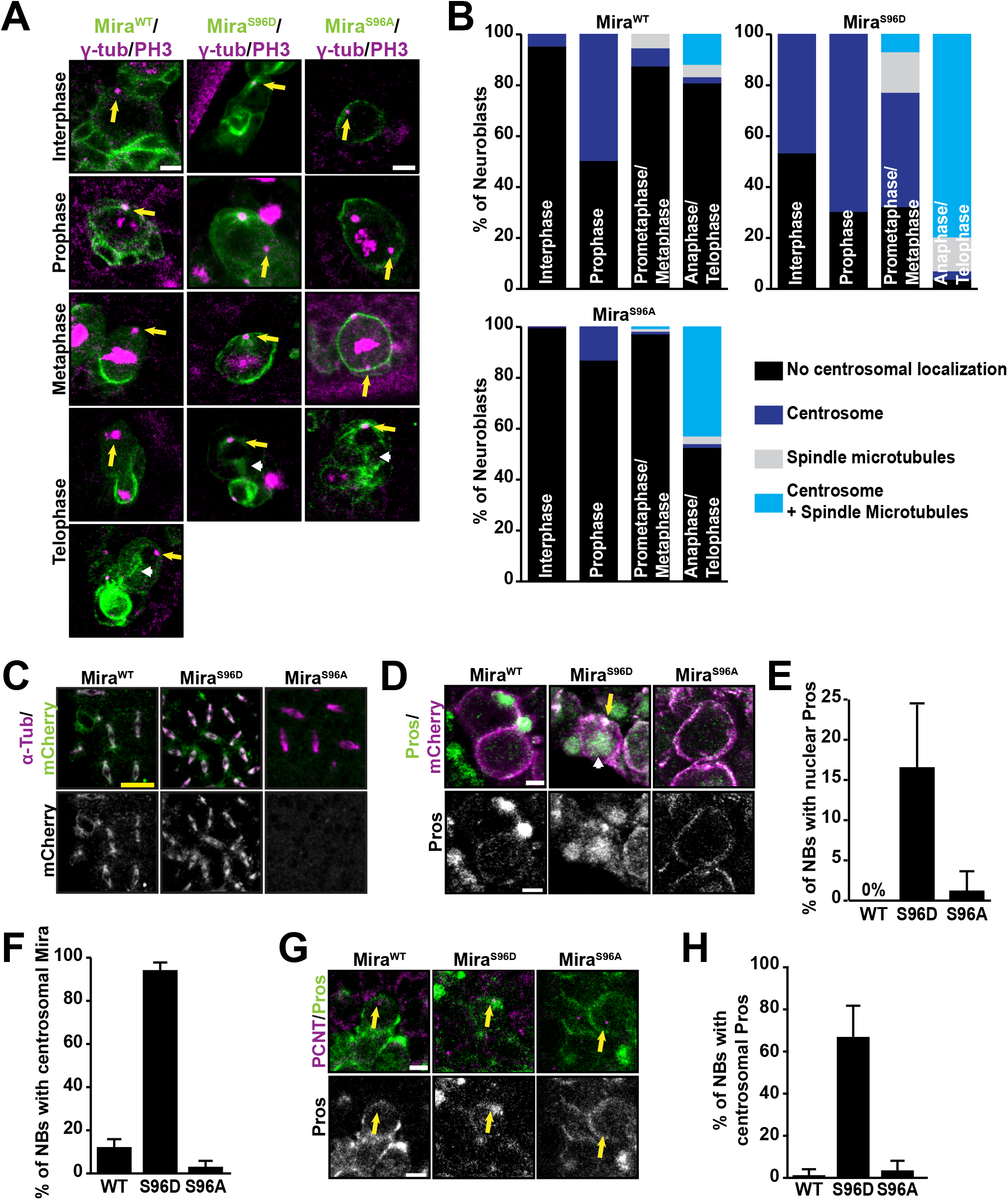
Phosphorylation of Mira residue S96 is required and sufficient for centrosomal/mitotic spindle localization. (**A**) Localization of Mira^WT^-mCherry, Mira^S96D^-mCherry, and Mira^S96A^-mCherry in neuroblasts. Mira alleles were visualized with DsRed antibody to distinguish them from endogenous Mira. Arrows point to centrosomes and arrowheads indicate spindle microtubules. Scale bars, 5 μm. (**B**) Quantification of centrosomal/spindle localization of Mira^WT^, Mira^S96D^, and Mira^S96A^ throughout the cell cycle. Interphase, n=119 NBs (Mira^WT^), 151 (Mira^S96D^), 135 (Mira^S96A^); Prophase, n=50 (Mira^WT^), 30 (Mira^S96D^), 52 (Mira^S96A^); Prometaphase/Metaphase, n=70 (Mira^WT^), 52 (Mira^S96D^), 94 (Mira^S96A^); Anaphase/Telophase, n=41 (Mira^WT^), 30 (Mira^S96D^), 67 (Mira^S96A^). (**C**) Localization of Mira^WT^, Mira^S96D^, and Mira^S96A^ in syncytial embryos. Scale bar, 25 μm. (**D**) Localization of Pros in embryonic neuroblasts homozygous for Mira^WT^, Mira^S96D^, and Mira^S96A^. Arrow indicates centrosome and arrowhead indicates nuclear Pros. Scale bar, 5 μm. (**E**) Quantification of nuclear localization of Pros in embryonic neuroblasts homozygous for Mira^WT^ (n=11 embryos), Mira^S96D^ (n=10), and Mira^S96A^ (n=8). Error bars, 1 SD. (**F**) Quantification of centrosomal Mira localization in Mira^WT^ (n=9 embryos), Mira^S96D^ (n=9), and Mira^S96A^ (n=6) homozygous mutant embryonic neuroblasts. Error bars, 1 SD. (**G**) Localization of Pros to the centrosome (marked by PCNT staining) in embryonic neuroblasts homozygous for Mira^WT^, Mira^S96D^, and Mira^S96A^. Arrow indicates centrosome. Scale bar, 5 μm. (**H**) Quantification of centrosomal Pros localization in Mira^WT^ (n=9 embryos), Mira^S96D^ (n=9), and Mira^S96A^ (n=9) homozygous embryonic neuroblasts. Error bars, 1 SD.

As the lack of centrosomal localization of Mira^S96A^ could be due to sequestration at the cortex rather than an inability to localize to the centrosome, we examined the localization of these alleles in syncytial embryos as syncytial embryos do not have a cell membrane and Mira has been shown to localize to the mitotic spindle in syncytial embryos (Mollinari *et al*, 2002). We observed that both Mira^WT^ and Mira^S96D^ localized to the mitotic spindle as demonstrated by colocalization with α-tubulin, whereas Mira^S96A^ did not (Fig. 5C), indicating that reduction of Mira^S96A^ localization to the centrosome/spindle is not due to sequestration at the cortex and that phosphorylation of S96 is required for localization of Mira to the centrosome/spindle.

PP4 has also been reported to be required for dephosphorylation of T591, however, we did not observe an increase in centrosomal localization when a phosphomimetic construct, Mira^T591D^, was expressed in neuroblasts, indicating T591 phosphorylation does not contribute to this phenotype (Figs. S4A,B). Therefore, the mislocalization of Mira to the centrosome observed in Kin17 knockdown neuroblasts is likely due to phosphorylation of Mira on S96.

Next, we asked whether mislocalization of Mira to the centrosome caused by the phosphorylation of S96 is responsible for the aberrant localization of Pros in the nucleus and the centrosome observed in Kin17 knockdown neuroblasts. We did not observe a phenotype in larval brains heterozygous for the S96 mutant alleles potentially due to the presence of endogenous wild-type Mira. Therefore, we examined Pros localization in stage 9-11 homozygous mutant embryonic neuroblasts. We found that Pros localized primarily to the cortex in interphase *mira*^*WT*^ and *mira*^*S96A*^ neuroblasts. However, we observed that Pros localized to the nucleus at a rate of 16.5% in *mira*^*S96D*^ neuroblasts compared to 0% and 1.2% for *mira*^*WT*^ and *mira*^*S96A*^ neuroblasts respectively (Figs. 5D,E). At interphase, Mira^S96D^ localized to the centrosome in 94% of neuroblasts, whereas Mira^WT^ and Mira^S96A^ localized to the centrosome in 12% and 2.8% of neuroblasts, respectively (Figs. 5D,F). Accordingly, we observed that Pros localized to the centrosome in about 66.7% of interphase *mira*^*S96D*^ neuroblasts (Figs. 5G,H), but in less than 3% of *mira*^*WT*^ or *mira*^*S96A*^ neuroblasts. These results suggest that nuclear localization of Pros in Kin17 knockdown neuroblasts is likely due to phosphorylation of Mira at S96 and subsequent mislocalization of Mira to the centrosome.

### Reduction of Protein Phosphatase 4 levels leads to the localization of Mira to the centrosome

Aberrant Mira phosphorylation could be due to an increase in aPKC activity or a decrease in phosphatase activity. We first investigated if increased aPKC activity accounts for the aberrant Mira localization in Kin17 knockdown neuroblasts. We observed that localization of the apical components Bazooka and aPKC were not altered in Kin17 knockdown (Figs. S5A,B) and reduction of aPKC levels in the *Kin17 RNAi* background utilizing the null allele *aPKC*^*K06403*^ did not rescue brain size, loss of Pros+ neurons, or nuclear Pros localization (Figs. S5C,D), suggesting that Kin17 may function through a phosphatase.

Falafel (Flfl), the targeting subunit of Protein Phosphatase 4 (PP4), is required for Mira localization to the basal domain and interacts directly with Mira, and PP4 is known to be required for dephosphorylation of Mira during interphase(Sousa-Nunes *et al*, 2009; Zhang *et al*, 2016). We tested the role of PP4 by utilizing RNAi to the catalytic subunit PP4-19c and Flfl. We found that both *PP4-19c* and *Flfl RNAi* led to an increase of Mira localization to the centrosome through the cell cycle in a pattern similar to that observed in *Kin17 RNAi*. PP4-19c knockdown leads to localization of Mira to the centrosome in 45.0% of interphase neuroblasts, 78.1% of prophase neuroblasts, 63.0% of prometaphase/metaphase neuroblasts, and 21.9% of anaphase/telophase neuroblasts (Figs. 6A,C). Flfl knockdown led to a similar pattern with Mira localizing to the centrosome in 37.5% of interphase neuroblasts, 74.5% of prophase neuroblasts, 62.4% of prometaphase/metaphase neuroblasts, and 31.3% of anaphase/telophase neuroblasts. (Figs. 6B,C). However, PP4-19c or Flfl knockdown did not affect the preferential localization of Mira to the apical centrosome or cortical localization (Figs. S6A,B). Nor did we observe nuclear Pros localization or reduction in the mitotic rate of neuroblasts. To determine if the lack of nuclear Pros and other phenotypes was due to incomplete knockdown, we examined *flfl*^*n42*^ mutant neuroblasts. We found that *flfl*^*n42*^ mutants exhibited nuclear Pros localization in 52.5% of neuroblasts and a significant reduction in the mitotic rate (Figs. S6C-G). Taken together, the accumulation of Mira at the centrosome/spindle in PP4-19C and Flfl knockdown neuroblasts supports the hypothesis that decreased phosphatase activity leads to accumulation of Mira at the centrosome in Kin17 knockdown.

**Figure 6.**
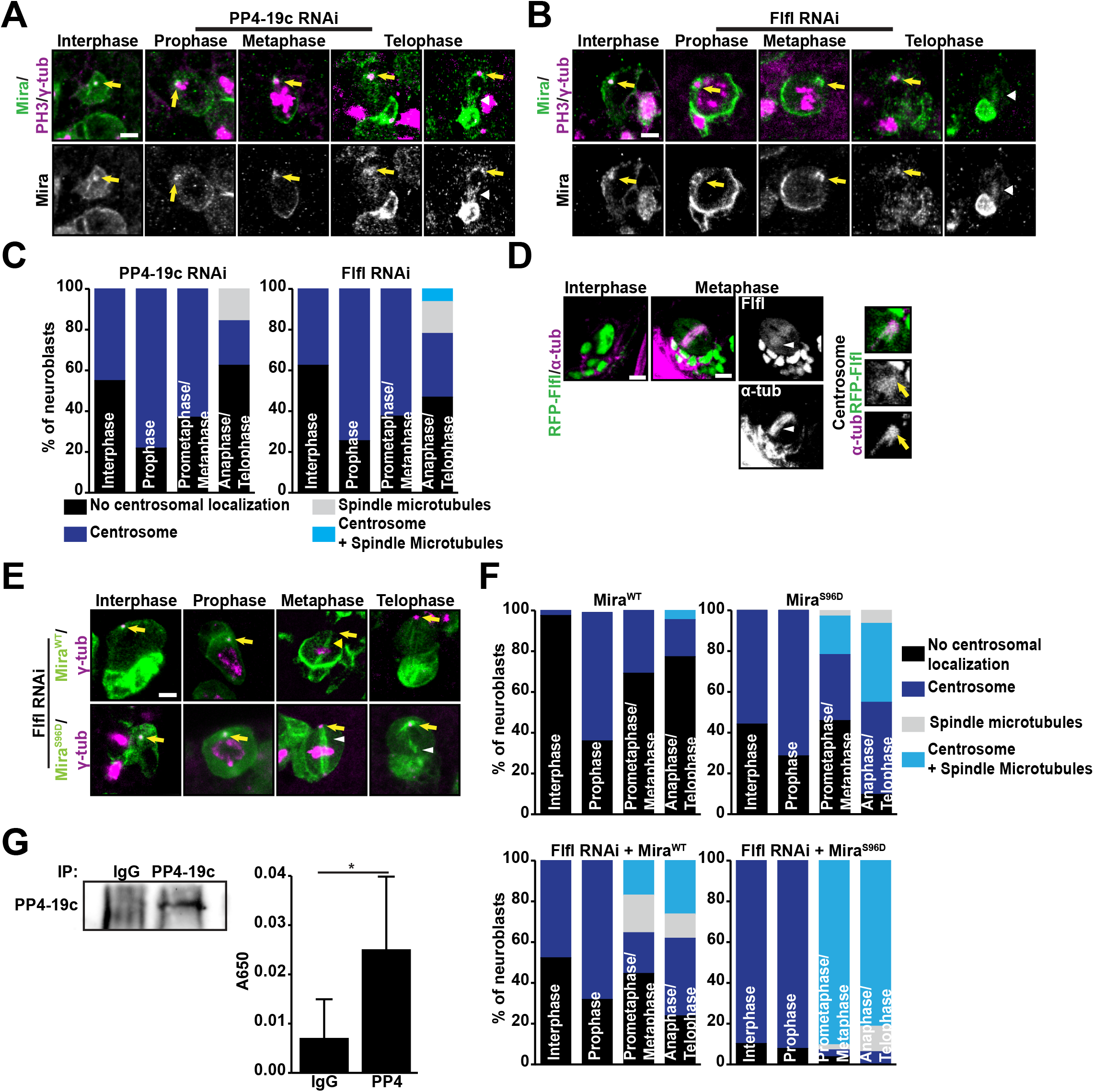
PP4 is required for displacement of Mira from the centrosome. (**A**) Localization of Mira in *PP4-19c RNAi* neuroblasts throughout the cell cycle. Arrows point to centrosomes and arrowheads indicate spindle microtubule localization of Mira. Scale bar, 5 μm. (**B**) Localization of Mira in *Flfl RNAi* neuroblasts throughout the cell cycle. Arrows point to centrosomes and arrowheads indicate spindle microtubule localization of Mira. Scale bar, 5 μm. (**C**) Quantification of Mira localization during the cell cycle in *PP4-19c* and *Flfl RNAi* neuroblasts. Interphase, n=200 NBs (PP4-19c), 136 (Flfl); Prophase, n=32 (PP4-19c), 98 (Flfl); Prometaphase/Metaphase, n=81 (PP4-19c), 132 (Flfl); Anaphase/Telophase, n=32 (PP4-19c), 32 (Flfl). (**D**) Localization of RFP-Flfl in wild-type neuroblasts. Arrowheads indicate the mitotic spindle and arrows point to the centrosome. Scale bar, 5 μm. (**E**) Mira^WT^ and Mira^S96D^ localization in Flfl knockdown neuroblasts. Arrows point to centrosomes and arrowheads indicate spindle microtubule localization of Mira. Scale bars, 5 μm. (**F**) Quantification of Mira^WT^ and Mira^S96D^ localization in Flfl knockdown neuroblasts. Interphase, n=40 NBs (Mira^WT^), 52 (Mira^S96D^), 130 (Mira^WT^, *Flfl RNAi*), 117 (Mira^S96D^, *Flfl RNAi*); Prophase, n=25 (Mira^WT^), 28 (Mira^S96D^), 69 (Mira^WT^, *Flfl RNAi*), 51 (Mira^S96D^, *Flfl RNAi*); Prometaphase/Metaphase, n=26 (Mira^WT^), 37 (Mira^S96D^), 65 (Mira^WT^, *Flfl RNAi*), 53 (Mira^S96D^, *Flfl RNAi*); Anaphase/Telophase, n=21 (Mira^WT^), 31 (Mira^S96D^), 42 (Mira^WT^, *Flfl RNAi*), 32 (Mira^S96D^, *Flfl RNAi*). (**G**) Dephosphorylation of Mira pS96 synthetic peptide. Error bars, 1 SD; *, p=0.03; n=4.

### Flfl localizes to the mitotic spindle

We next wanted to identify the localization of the PP4 complex in neuroblasts. In neuroblasts, Flfl has been previously reported to localize to the nucleus during interphase but with no specific localization during mitosis. However, in *Drosophila* D.mel-2 cells, Flfl has been reported to localize to the kinetochores where it forms a direct interaction with the kinetochore protein, CENP-C (Lipinszki *et al*, 2015), and PP4 is a centrosomal protein complex required for microtubule assembly in *Drosophila embryos* (Helps *et al*, 1998; Ou & Rattner, 2004). To visualize the localization of Flfl in neuroblasts, we utilized a tagged UAS-construct, *UAS-RFP-Flfl* (Sousa-Nunes *et al*, 2009). We found that during interphase, RFP-Flfl localized to the nucleus, while during mitosis we observed enrichment of RFP-Flfl at the centrosome and mitotic spindle in 67.3% of neuroblasts (Fig. 6D). The localization of Flfl to the centrosome/spindle is consistent with a role for PP4 in preventing mislocalization of Mira to the centrosome/spindle.

### PP4 dephosphorylates Mira at Serine-96

PP4 has not been shown to dephosphorylate Mira at S96. Thus we first performed a genetic interaction test to see if the *Flfl RNAi* phenotypes would be enhanced in the *mira*^*S96D*^ heterozygous background. We found that localization of Mira^WT^ during Flfl RNAi was like that of endogenous Mira, however, *Flfl RNAi* led to significant enhancement of the centrosomal/spindle localization of Mira^S96D^ throughout the cell cycle, with ∼90% of neuroblasts exhibiting centrosomal localization of Mira^S96D^ (Figs. 6E,F). We also observed increased defects in Mira^S96D^ cortical localization (Figs. S6G,H), supporting the hypothesis that PP4 dephosphorylates S96.

As these results do not demonstrate direct dephosphorylation of Miranda, we utilized an *in vitro* malachite green-based phosphatase assay. We immunoprecipitated the PP4 complex from *Drosophila* S2 cells (Fig. 6G, left) and incubated this with a synthetic phospho-peptide corresponding to the region surrounding S96 (92-FRTP(pSer)LPQR-100). We found that incubating PP4 immunoprecipitates with the Mira pS96 peptide led to a 3.6-fold increase in dephosphorylation compared to IgG immunoprecipitates (Fig. 6G, right), indicating that PP4 can directly dephosphorylate Miranda at S96.

### Flfl expression partially rescues the Kin17 phenotype

To determine if Kin17 functions upstream of Flfl, we performed rescue experiments by expressing RFP-Flfl in *Kin17 RNAi* neuroblasts. We first quantified the localization of Mira in neuroblasts overexpressing RFP-Flfl alone to ensure that it does not lead to a phenotype. We found that Mira localization during overexpression of RFP-Flfl was similar to that observed in control neuroblasts with Mira localized to the centrosome in 5.9% of interphase neuroblasts, 53.2% of prophase neuroblasts, 35.4% of prometaphase/metaphase neuroblasts, and 9.7% of anaphase/telophase neuroblasts (Figs. 7A,C). We then expressed RFP-Flfl in *Kin17 RNAi* neuroblasts. We found that expression of RFP-Flfl was able to partially rescue the phenotypes observed in *Kin17 RNAi* knockdown. We found that when RFP-Flfl was expressed in *Kin17 RNAi* neuroblasts, mis-localization of Mira to the centrosome was significantly reduced at all different phases of the cell cycle except prophase. Mira localized to the centrosome in 39.1% of interphase neuroblasts, 58.8% of prophase neuroblasts, 43.4% of metaphase neuroblasts, and 29.4% of telophase neuroblasts (Figs. 7B,C). Relative to controls and *Kin17 RNAi*, this corresponds to a reduction of aberrant centrosomal localization by about half. In addition, there was a slight decrease in cortical localization defects, although localization to the basal centrosome was not improved (Figs. S7A-C).

**Figure 7.**
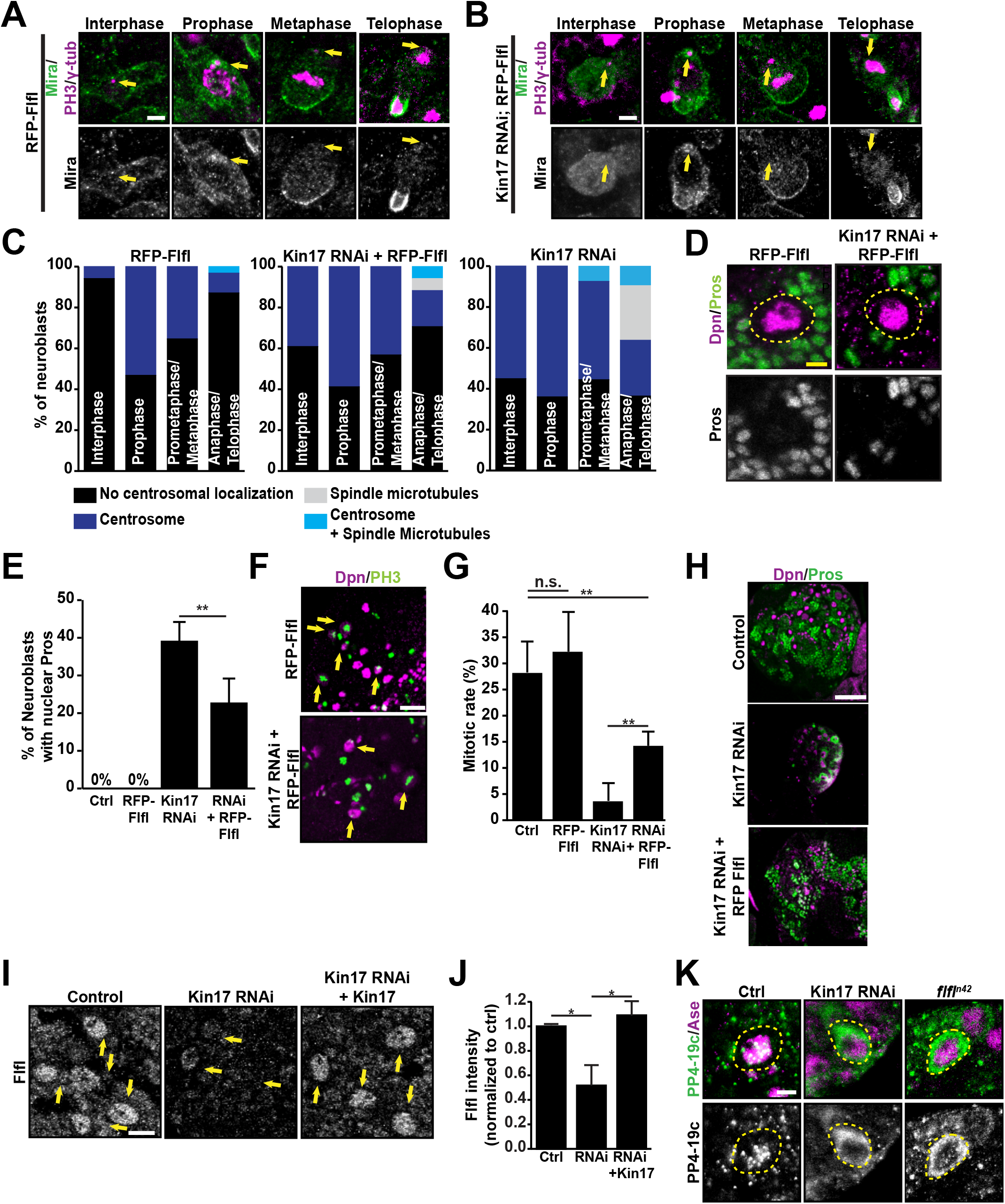
Flfl expression partially rescues *Kin17 RNAi* phenotypes. (**A**) Localization of Mira in neuroblasts overexpressing RFP-Flfl similar to that in wild-type neuroblasts throughout the cell cycle. Arrows indicate centrosomes. Scale bar, 5 μm. (**B**) Overexpressing RFP-Flfl partially rescues aberrant centrosomal localization of Mira in *Kin17 RNAi* neuroblasts throughout the cell cycle. Arrows indicate centrosomes. Scale bar, 5 μm. (**C**) Quantification of Mira localization in neuroblasts overexpressing RFP-Flfl in the control or *Kin17 RNAi* background throughout the cell cycle. Interphase, n=102 NBs (Flfl), 46 (*Kin17 RNAi*; Flfl), 49 (*Kin17 RNAi*); Prophase, n=62 (Flfl), 34 (*Kin17 RNAi;* Flfl), 25 (*Kin17 RNAi*); Prometaphase/Metaphase, n=65 (Flfl), 37 (*Kin17 RNAi;* Flfl), 27 (*Kin17 RNAi*); Anaphase/Telophase, n=31 (Flfl), 17 (*Kin17 RNAi*; Flfl), 11 (Kin17 RNAi). (**D**) Overexpressing RFP-Flfl partially rescues aberrant nuclear localization of Pros in *Kin17 RNAi* neuroblasts. Dotted lines indicate neuroblasts. Scale bars, 5 μm. (**E**) Quantification of Pros localization to the nucleus in control (n=10 brains), *Kin17 RNAi* (n=9), RFP-Flfl (n=5), and *Kin17 RNAi*; RFP-Flfl (n=13) brains. **, *p*<0.01. (**F**) pH3 staining in neuroblasts expressing RFP-Flfl in the control and *Kin17 RNAi* background. Arrows indicate mitotic neuroblasts. Scale bars, 20 μm. (**G**) Quantification of mitotic rate in neuroblasts expressing RFP-Flfl in the control and *Kin17 RNAi* background. Error bars, 1 SD, **, *p*<0.01; n.s, not significant; n=11 brains for all conditions. (**H**) Pros^+^ neurons are partially rescued by expression of RFP-Flfl in *Kin17 RNAi* larval brains. Scale bar, 50 μm. (**I**) Flfl staining in control, *Kin17 RNAi*, and Kin17 rescue neuroblasts. Arrows indicate neuroblasts (identified by size and CD8-GFP expression (not shown)). Scale bar, 5 μm. (**J**) Quantification of relative Flfl staining intensities in control, *Kin17 RNAi*, and Kin17 rescue neuroblasts. Error bars, 1SD; *, *p*<0.05, n=3 replicates; Control (n=5 brains, 5, 7); Kin17 RNAi (n=3,5,6); *Kin17 RNAi* + Kin17 (n=5,5,5). (**K**) PP4-19c localization in control, *Kin17 RNAi, Kin17 RNAi* + RFP-Flfl, and *flfl*^*n42*^ neuroblasts. Dotted lines indicate neuroblasts. Scale bar, 5 μm.

Consistent with the partial rescue of the mislocalization of Mira to the centrosome, the number of neuroblasts expressing nuclear Pros was significantly reduced from 39.1 ± 5.3% in *Kin17 RNAi* to 22.7 ± 6.8% (Figs. 7D,E) after expressing RFP-Flfl. Accordingly, the mitotic rate was significantly increased from 3.5 ± 3.4% in *Kin17 RNAi* neuroblasts to 14.7 ± 2.7% (Figs. 7F,G). The Pros^+^ neurons, which were largely missing in *Kin17 RNAi* brains, were also partially rescued by the expression of RFP-Flfl (Fig. 7H). Together these data show that over-expression of Flfl can partially rescue the phenotype generated by *Kin17 RNAi*, indicating that Flfl likely functions downstream of Kin17.

### Kin17 regulates cellular levels of Flfl

We then examined the expression of Flfl and PP4-19c in *Kin17 RNAi* neuroblasts. We quantified nuclear Flfl levels during interphase as Flfl is known to localize to the nucleus of interphase neuroblasts (Sousa-Nunes *et al*, 2009) and would provide a consistent readout of Flfl levels. We found that Kin17 RNAi led to a reduction in Flfl levels to about half that observed in control neuroblasts. Additionally, we found a change in PP4-19c localization from nuclear to cytoplasmic at interphase (Figs. 7I-K). A previous study showed that in *Dictyostelium*, SMEK, a homolog of Flfl, is required for PP4C to localize to the nucleus (Mendoza *et al*, 2007). Therefore, the cytoplasmic localization of PP4-19c in *Kin17 RNAi* neuroblasts could be due to the reduction in Flfl expression. Indeed, PP4-19c became cytoplasmic in *flfl*^*n42*^ mutant neuroblasts also (Fig. 7K) and Flfl expression in *Kin17 RNAi* neuroblasts rescues PP4-19c localization (Figs. S7D,E).

### The minor spliceosome component U6atac regulates Mira localization and Flfl expression

As previous experiments suggest that Kin17 may be involved in splicing and we found that Kin17 led to a reduction in the levels of Flfl in neuroblasts, we hypothesized that Kin17 may be involved in the splicing of *the flfl* transcript. To test this hypothesis, we first looked at the phenotypes of Mira localization in mutants for components of both the major and the minor spliceosome. Knockdown of the major spliceosome component, U2A, did not lead to Mira mislocalization (Fig. S8A,B).

However, a mutant allele for *u6atac* (*u6atac*^*k01105*^), a minor spliceosome component, led to mislocalization of Mira similar to that observed in *Kin17 RNAi*. Mira localized to the centrosome in 69.6%, 76.1%, 51.5%, and 52.6% of *u6atac*^*k01105*^ neuroblasts at interphase, prophase, metaphase, and anaphase/telophase, respectively (Figs. 8A,B). In addition to the overall increase in centrosomal localization, localization of Mira to the basal centrosome was also increased at prophase and prometaphase/metaphase (Fig. S8C). Furthermore, we observed an increase in cytoplasmic localization and a reduction in cortical localization of Mira in *u6atac*^*k01105*^ mutants (Figs. S8D,E).

**Figure 8.**
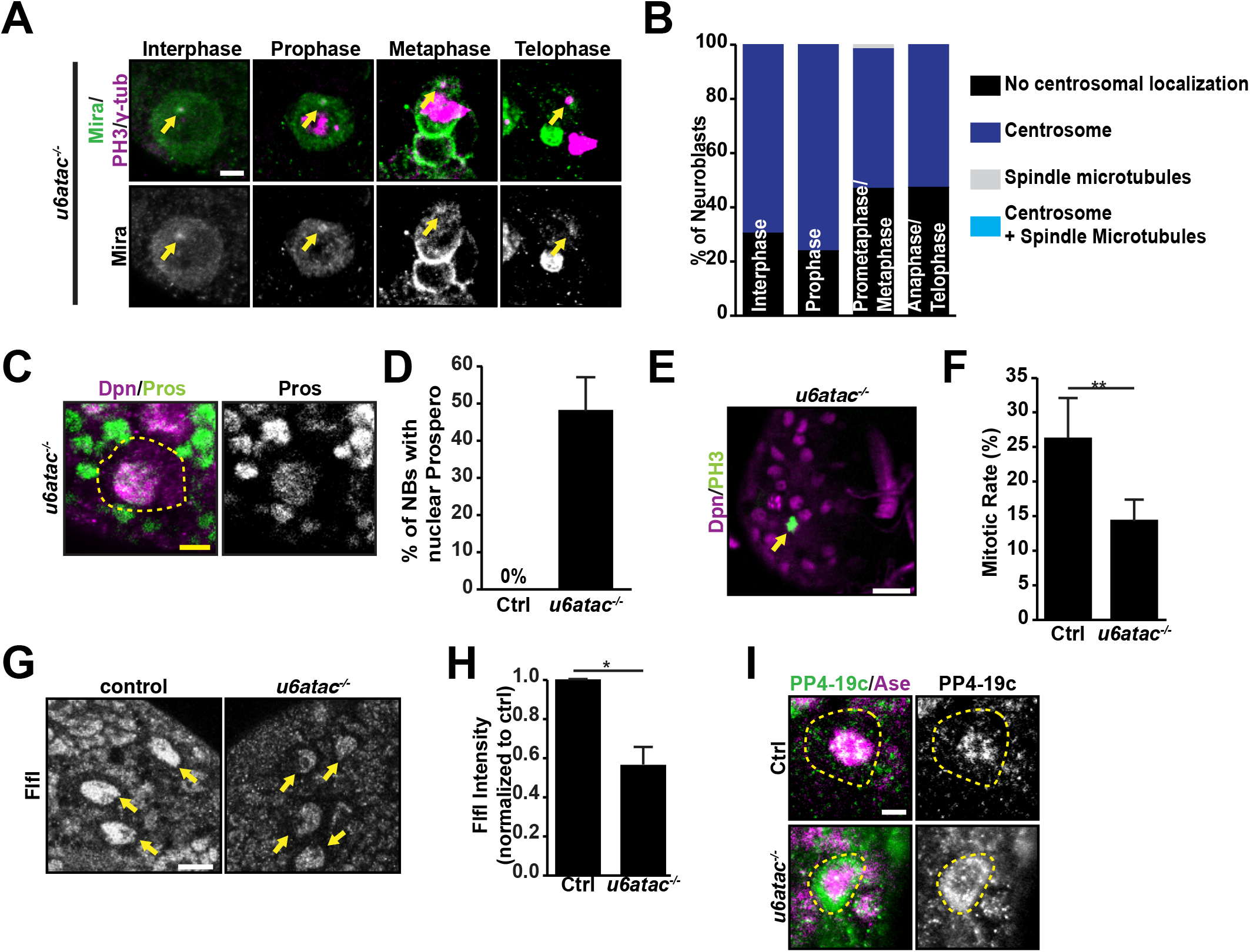
The minor spliceosome component U6atac regulates Mira localization and Flfl levels. (**A**) Localization of Mira throughout the cell cycle in *u6atac*^*K01105*^ (*u6atac*^*-/-*^) mutant neuroblasts. Arrows point to centrosomes. Scale bar, 5 μm. (**B**) Quantification of Mira centrosomal localization in *u6atac*^*-/-*^ mutant neuroblasts. n=69 NBs (Interphase), 46 (Prophase), 66 (Prometaphase/Metaphase), 19 (Anaphase/Telophase). (**C**) Nuclear localization of Pros in *u6atac*^*-/-*^ mutant neuroblasts. Dotted lines indicate neuroblasts. Scale bar, 5 μm. (**D**) Quantification of nuclear Pros localization in control (n=6 brains) and *u6atac*^*-/-*^ (n=10) neuroblasts. (**E**) PH3 staining in *u6atac*^*-/-*^ mutant brains. Arrows indicate mitotic cells. Scale bar, 20 μm. (**F**) Quantification of mitotic rate in control (n=13) and *u6atac*^*-/-*^ (n=7) brains. Error bars, 1 SD; **, *p*<0.01. (**G**) Flfl staining in *u6atac*^*-/-*^ mutant neuroblasts (arrows). Scale bar, 10 μm. (**H**) Quantification of relative Flfl staining intensities in control and *u6atac*^*-/-*^ mutant neuroblasts. Error bar, 1 SD; *, *p*<0.05.; n=3 replicates of 5 brains for each condition. (**I**) PP4-19c staining in *u6atac*^*-/-*^ mutant neuroblasts. Dotted lines indicate neuroblasts. Scale bar, 5 μm.

Consistent with the mislocalization of Mira, Pros mislocalized to the nucleus in 48.1 ± 9.2% of *u6atac*^*k01105*^ mutant neuroblasts (Figs. 8C,D). Consequently, the mitotic rate of *u6atac*^*k01105*^ mutant neuroblasts was reduced to 14.4 ± 3.1% from 26.3 ± 5.9% in the wild-type control (Figs. 8E,F). Additionally, loss of U6atac also led to a reduction in Flfl levels to ∼44% that of control and mislocalization of PP4-19c to the cytoplasm (Figs. 8G-I). These results indicate that loss of U6atac leads to phenotypes similar to that observed in Kin17 knockdown, suggesting that Kin17 could function in splicing of the *flfl* pre-mRNA.

### Kin17 and the minor spliceosome are required for the splicing of Flfl

Consistent with a potential role in splicing, Kin17-HA localized to the nucleus when expressed in neuroblasts (Fig. 9A). To assess if Kin17 and U6atac are involved in splicing the *flfl* pre-mRNA, we performed reverse transcriptase PCR utilizing primers that span introns and observed two products corresponding to the *flfl* pre-mRNA and mRNA. For *Kin17 RNAi* and *u6atac*^*k01105*^, we observed a higher abundance of the pre-mRNA than the spliced mRNA when compared to control (Fig. 9B) and this was observed for several different introns within the *flfl* pre-mRNA. To quantify this, we performed quantitative real-time PCR using primers that distinguish between the pre-mRNA and mRNA. We found that in both *Kin17 RNAi* and *u6atac*^*k01105*^ larval brains, the spliced form of *flfl* was reduced, whereas levels of the pre-mRNA were increased (Fig. 9C). These data indicate that reduction of Kin17 and U6atac levels lead to a decrease in the amount of spliced transcript and an increase in the amount of pre-mRNA, indicating that Kin17 and the minor spliceosome are required for the splicing of *flfl* mRNAs. Furthermore, immunoprecipitation of Kin17-HA from embryonic neuroblasts showed that Kin17 preferentially interacts with the *flfl* pre-mRNA (Figure 9ßD). Together these data support the hypothesis that Kin17 is required for proper splicing of the *flfl* mRNA and that this leads to a reduction in Flfl protein levels/activity and defects in cortical localization of Mira.

**Figure 9.**
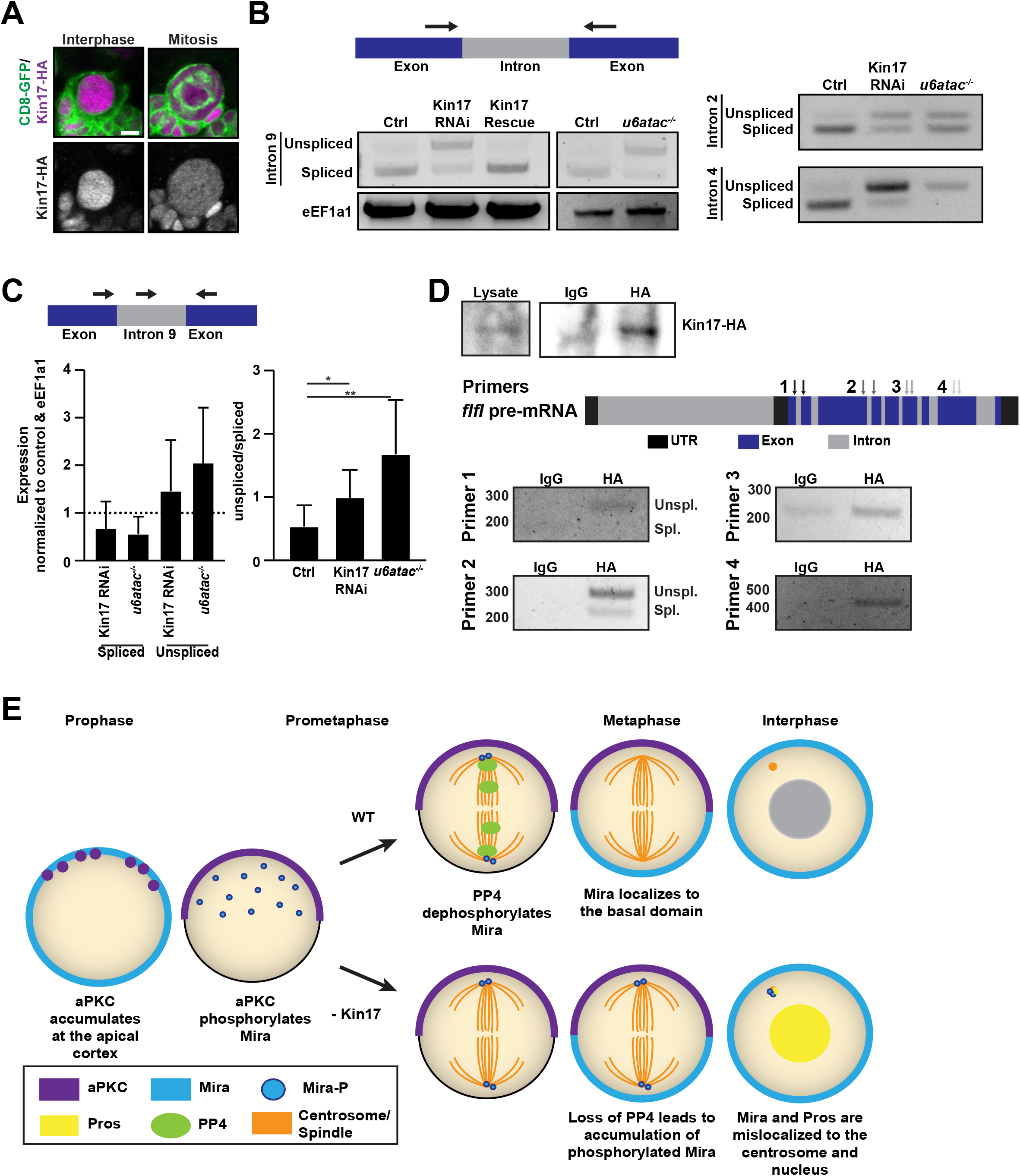
Regulation of the splicing of Flfl transcripts by Kin17 and U6atac and a proposed working model of Kin17. (**A**) Localization of Kin17-HA in neuroblasts during interphase and mitosis. Scale bar, 5 μm. (**B**) RT-PCR for Flfl spliced and unspliced forms from RNAs isolated from control, *Kin17 RNAi*, Kin17 rescue, and *u6atac*^*-/-*^ brains. The schematic at the top shows a part of the flfl gene structure and location of primers (arrows). (**C**) Quantitative PCR results for Flfl spliced and unspliced isoforms displayed as ΔΔCt and RIV. n=8; Error bars, 1 SD; *, *p*<0.05; \*\**p*<0.01. The top panel shows position of the primers. (**D**) RNA immunoprecipitation of *flfl* mRNA by Kin17-HA in *Drosophila* embryos. Top panel demonstrates IP of the Kin17-HA construct. Middle panel shows a schematic of *flfl* pre-mRNA and the position of RIP Primers. Bottom panels show PCR products obtained from cDNA libraries generated from the Kin17-HA and IgG immunoprecipitates. (**E**) A proposed working model. During prophase, aPKC accumulates at the cell cortex where it phosphorylates Mira at Serine-96. This phosphorylation event leads to Mira displacement to the cytoplasm where it accumulates on the centrosome during Prophase. At the centrosome, PP4 dephosphorylates Mira allowing it to dissociate from the centrosome and localize to the basal domain. However, in Kin17 knockdown, PP4 activity is reduced, which results in defects in Mira dephosphorylation and aberrant accumulation at the centrosome. Subsequently, Pros also accumulates at the centrosome and mislocalizes to the nucleus.

## Discussion

We have identified a novel factor, Kin17, which is essential for the proper localization of Mira in *Drosophila* neuroblasts. The localization of Mira during asymmetric cell division is essential to the proper segregation of the fate determinants to the daughter cell and the regulation of its phosphorylation state via an interplay between kinases and phosphatases is essential to this localization. We demonstrate that Kin17 regulates the levels of Flfl, a PP4 subunit, by regulating the splicing of its transcript. PP4 is likely required for the dephosphorylation of Mira at S96 at the centrosome/mitotic spindle beginning in prophase and completing dephosphorylation by the end of mitosis. Dephosphorylation of Mira prevents its accumulation in the cytoplasm and at the centrosome/mitotic spindle (Figure 10E). Knockdown of Kin17 leads to defects in splicing of *flfl* transcripts and reduction in Flfl expression, which in turn results in reduced PP4 activity and mislocalization of PP4. Consequently, Mira cannot be fully dephosphorylated at S96 after being phosphorylated by aPKC and displaced from the apical cortex at prophase/metaphase. The phosphorylated Mira accumulates in the cytoplasm and the centrosome/mitotic spindle instead of binding to the basal cortex (Figure 10E).

Kin17 has not been previously well-studied in *Drosophila*, however, it has been identified in mammalian systems as essential for DNA repair and replication. Kin17 has also been shown to interact with the spliceosome through mass spectrometry but this role has not been investigated further previously (Herold *et al*, 2009; Rappsilber *et al*, 2002; Gaspar *et al*, 2021), nor has it been determined whether Kin17 functions as a general splicing factor or regulates the splicing of specific transcripts. Here we show that Kin17 is required for the proper splicing of Flfl in neuroblasts and Kin17 knockdown phenocopies mutants for the minor spliceosome component *u6atac*, demonstrating that Kin17 indeed functions to regulate splicing of transcripts. In the mass spectrometry analysis, Kin17 was only found in complexes isolated from both HeLa and *Drosophila* Kc cells using *fushi tarazu* as bait but not *zeste* (Herold *et al*, 2009), suggesting that Kin17 likely regulates splicing of specific transcripts rather than acting as a general minor spliceosome factor. Otherwise, one would expect Kin17 to be required for splicing of other transcripts such as *pros* transcripts, which are spliced by both the major and minor spliceosomes in neuroblasts (Otake *et al*, 2002), and Kin17 knockdown would lead to a reduction in Pros expression. Yet, our results show the *Kin17 RNAi* phenotypes were partially rescued rather than exacerbated in *pros*^*17/+*^ heterozygous background, suggesting that Pros expression is not reduced in *Kin17 RNAi* neuroblasts and Kin17 unlikely regulates splicing of *pros* transcripts. Therefore, our work demonstrates that Kin17 functions as a splicing factor in neuroblasts to regulate splicing of specific transcripts and proper localization of cell fate determinants. These findings could be potentially very helpful for further investigating underlying mechanisms of pathogenesis of various cancers, including breast, liver, colorectal, cervical, and non-small cell lung cancers that have increased levels of Kin17 expression (Zeng *et al*, 2011; Zhang *et al*, 2017a, 2017b; Kou *et al*, 2014; Ruan *et al*, 2018).

Mira localization to the mitotic spindle/centrosome has been observed in the syncytial embryo (Mollinari *et al*, 2002), and interactions between Mira and the microtubules have also been observed in the anterior pole of oocytes (Irion *et al*, 2006). Mutants in polarity proteins can lead to localization of Mira to the centrosome and mitotic spindle in neuroblasts, raising the question of why Mira does not localize to the mitotic spindle in wild-type neuroblasts, and whether localization to the mitotic spindle during mitosis leads to consequences for asymmetric cell division. Our work provides evidence that localization to the centrosome and potentially the mitotic spindle (based on the S96 alleles) does occur in wild-type neuroblasts but it is likely transient and appears to be cell-cycle dependent. Our work also reveals that localization of Mira to the centrosome/spindle depends on the phosphorylation status of Mira at S96. However, unlike in wild-type neuroblasts that have Mira preferentially localized to the apical centrosome during the division, in Kin17 knockdown neuroblasts, such preference is less obvious. Instead, Mira localizes to both apical and basal centrosomes. Although the exact underlying mechanisms remains to be investigated, one potential explanation could be that in wild-type neuroblasts, the apical centrosome is closer to the apical cortex, and therefore, when Mira is phosphorylated, it will first encounter the apical centrosome where it will be dephosphorylated by PP4 and dissociated from the cortex.

Interestingly, we observed an increase of Mira localization to the centrosome in *PP4* and *Flfl RNAi* neuroblasts with a minimal increase in cytoplasmic Mira. This suggests that centrosomal/spindle localization of Mira may be more sensitive to changes in the levels of PP4 activity. The phosphorylated Mira may preferentially localize to the centrosome/spindle and bring its cargo protein Pros to the centrosome as well as indicated by the colocalization of Pros and Mira to the centrosome in *Kin17 RNAi* neuroblasts. However, how knockdown of Kin17 or reduction in PP4 activity leads to aberrant nuclear accumulation of Pros still needs to be investigated. *mira* null mutants exhibit nuclear Pros localization (Hannaford *et al*, 2018), suggesting that Pros must be tethered to something to prevent nuclear localization and that defects in S96 dephosphorylation causes tethering to the cortex to be impaired so Mira/Pros then localizes to the centrosome. It is tempting to speculate that when the amount of phosphorylated Mira exceeds the binding capacity of the centrosome/spindle, it may “overflow” to the cytoplasm leading to untethered Pros being transported into the nucleus.

Although PP4 has been implicated in dephosphorylation of Mira at T591 (Zhang *et al*, 2016), our results suggest that PP4 also dephosphorylates Mira at a novel site S96 and that T591 phosphorylation does not contribute to the observed phenotype. In fact, we observed a slight decrease in T591D centrosomal localization, when compared to controls. This may be due to the fact that T591 dephosphorylation is required prior to aPKC phosphorylation of Mira to ensure proper localization (Zhang *et al*, 2016). This indicates that Mira removal from the centrosome is specific to PP4 dephosphorylating S96.

In summary, our studies identify a novel factor, Kin17, that regulates the proper localization of Mira during asymmetric cell division through regulation of the splicing of *flfl* transcripts and provides evidence to demonstrate that dephosphorylation of Mira at S96 by PP4 is essential for preventing aberrant localization of Mira to the centrosome in neuroblasts.

## Methods

### Fly stocks

*insc-Gal4* (Betschinger *et al*, 2006) was used for transgene expression and *UAS-CD8-GFP* was used to mark neuroblast lineages. UAS transgenes for RNAi knockdown or overexpression include: *UAS-Kin17 RNAi* (Ni *et al*, 2009) (#55692; Bloomington Drosophila Stock Center (BDSC) (IN, USA)), *UAS-kin17* (generated for this work), *UAS-kin17 HA* (this work), *UAS-RFP-flfl* (Sousa-Nunes *et al*, 2009) (#66538, BDSC), *UAS-numb*^γ*CT*^*-GFP* (Song & Lu, 2012), *UAS-flfl RNAi* (Sousa-Nunes *et al*, 2009) (#66541, BDSC), *UAS-U2A-RNAi* (Ni *et al*, 2009) (#33671; BDSC), and *UAS-PP419c RNAi* (Ni *et al*, 2009) (#57823; BDSC). The Mira phosphomimetic alleles, *mira*^*WT*^*-HA-mCherry, mira*^*S96A*^*-HA-mCherry, mira*^*S96D*^*-HA-mCherry* (Hannaford *et al*, 2018) were generously provided by Jens Januschke. *snRNA:U6atac*^*k01105*^ (Spradling *et al*, 1999) (#10492, BDSC), *flfl*^*n42*^ (Sousa-Nunes *et al*, 2009) (#66534, BDSC), *pros*^*17*^ (Doe *et al*, 1991) (#5458, BDSC) and *aPKC*^*K06403*^ (Spradling *et al*, 1999) (#10622, BDSC) mutant alleles were used for phenotypic analysis. Mutant chromosomes were balanced over *CyO, WeeP* or *TM6B, Tb* for larval analysis or *TM3, Tw-Gal4, UAS-2xGFP* for embryonic analysis. Lines were raised at 25°C on standard fly food.

### RNAi knockdown and transgene analysis

For RNAi knockdown and transgene overexpression, embryos were collected for 24 hours at 25°C, staged at 25°C for an additional 24 hours, and then shifted to 29°C until dissection at third instar larval stages. To visualize RFP-Flfl localization, crosses were collected at 25°C and shifted to 29°C, 24 hrs prior to dissection at third instar larval stages. Crosses involving *UAS-numb*^γ*CT*^*-GFP* were performed at 25°C. For syncytial embryo staining, embryos were collected for 1 hour and staged for 1 hour at 25°C before fixation. For stage 9-11 embryonic neuroblast staining, embryos were collected for 4 hours and staged for 4 hours at 25°C before fixation.

#### Immunostaining and confocal microscopy

Third instar larval brains were dissected in PBS and fixed within 20 minutes in 4% formaldehyde (Fisher (NH, USA)) in PBS for 20 minutes except for brains stained for Mira which were fixed for 35 minutes. Following fixation, brains were blocked for 30 min in PBS supplemented with 5% normal donkey serum (Jackson Immunoresearch (PA, USA)) and 0.3% Triton X-100 (Fisher). Brains were incubated with primary antibodies overnight at 4°C and secondary antibodies for 2 hours at RT. Embryos were stained as previously described (Rothwell & Sullivan, 2000).

Primary antibodies used: Dpn (Rabbit (Rb); gift from Y.N. Jan (Bier *et al*, 1992); 1:500); Pros (Mouse (Ms); Developmental Studies Hybridoma Bank (IA, USA); 1:5 (cortical/centrosomal localization), 1:50 (nuclear localization)); Pericentrin (Rb; Abcam; 1:500); PH3 (Ms; Cell Signaling Technology; 1:1000); Elav (Rat (Rt); DHSB; 1:100), α-tub (Ms; Sigma (MO, USA); 1:1000), γ-tub (Ms; Sigma; 1:1000), Mira (Rb; gift from Y.N. Jan; 1:1000); HA (Rb; Invitrogen (CA, USA); 1:500), Flfl (Rt; Gift from Z. Lipinszki (Lipinszki *et al*, 2015); 1:500), Baz (Gp; Gift from C. Doe (Siller *et al*, 2006); 1:500), aPKC (Ms; Santa Cruz Biotechnology (TX, USA); 1:500), GFP (Chicken (Ch); Aves Labs (OR, USA); 1:500), PP4-19c (Rb; Proteintech: 1:000), and DsRed (Rb; Takara Bio (CA, USA); 1:500). Secondary antibodies used: Alexafluor-488 (Ch, Invitrogen, 1:500), Alexafluor-647 (Rb/Ms, Invitrogen, 1:500), and RRX (Rb, Rt, Ms; Jackson Immunoresearch; 1:500).

Coverslips were mounted using antifade reagent in glycerol (Invitrogen). Images were taken using a Zeiss LSM 780 (Zeiss, Germany) inverted confocal microscope using a 40x 1.4 numerical aperture oil immersion objective. Images were collected using Zen Software (Zeiss) and processed with Fiji (Schindelin *et al*, 2009).

### Construction of plasmids and generation of transgenic lines

For pUAST-Kin17 and pUAST-Kin17 HA, the open reading frame was amplified from a larval brain cDNA library using CloneAmp HiFi PCR Premix (Takara Bio). For the HA-tagged version, the HA tag was integrated into the primer at the c-terminus of Kin17. Primers used were: Fwd (5’GGAATTCATGGGTCGCGCCGAGGTA-3’); Fwd. HA (5’-ATCGATGAATTCATGGGTCGCGCCGAGGTAGGT-3’); Reverse (5’-GCTCTAGACTAGGCGCCATGTAGTTTAGATAT-3’); and Rev. HA (5’-ATCGATCTCGAGTTCCTAAGCGTAATCTGGAACATCGTAAGGGTAGGCGCCATGTAGTTTAGATAT-3’). The PCR products were digested by EcoRI and XbaI or XhoI before cloning into the pUAST vector (Brand & Perrimon, 1993). UAS-Kin17 and UAS-Kin17 HA were integrated randomly into the genome of *w*^*1118*^ embryos and positive lines were mapped. Injections were performed by The Best Gene (CA, USA).

To generate pUAST-Mira T591 mutants, Mira was amplified from genomic DNA using the following primers: Mira Fwd (5’-TTCAGTAGATCTATGTCTTTCTCCAAGGCCAAG-3’) and Mira Rev (5’-TTCAGTCTCGAGGATGTTGCGCGCCTTGAGCAC-3’). Mira was cloned into pJET using the CloneJet PCR cloning kit (ThermoFisher) and T591A/D mutations were induced using quickchange PCR with the following primers: T591D Fwd (5’-CTGAGATCCTCCTCCCAGGATCTGCAGAGCGAGGTATCG-3’), T591D Rev (5’-CGATACCTCGCTCTGCAGATCCTGGGAGGAGGATCTCAG-3’), T591A Fwd (5’-CTGAGATCCTCCTCCCAGGCCCTGCAGAGCGAGGTATCG-3’), and T591 Rev (5’-CGATACCTCGCTCTGCAGGGCCTGGGAGGAGGATCTCAG-3’). Mira^WT^, Mira^T591A^, and Mira^T591D^ were digested from pJET using BglII and XhoI and cloned into pUAST-GFP (see above). UAS-Mira^WT^-GFP, UAS-Mira^T591D^-GFP, and UAS-Mira^T591A^-GFP were integrated randomly into the genome of *w*^*1118*^ embryos and positive lines were mapped. Injections were performed by Rainbow Transgenic Flies, Inc. (CA, USA).

### Cell Culture

*Drosophila* S2 cells were grown in Schneider’s medium (ThermoFisher Scientific) supplemented with 10% FBS (ThermoFisher Scientific) and 5% streptomycin/penicillin (Gibco) at 37°C.

### Phosphatase assays

S2 cells were lysed in lysis buffer (50 mM Tris-HCl pH 7.4, 50 mM NaCl, 1 mM EDTA, 0.1% NP40) supplemented with HALT protease inhibitor cocktail (Thermo Scientific (MA, USA)). Endogenous PP4 was immunoprecipitated with Protein A/G Plus Agarose beads (Santa Cruz Biotechnology) and 4 ug/mg protein of anti-PP4-19c antibody (Rb, Proteintech (IL, USA)) with anti-IgG (Rb, Invitrogen) as a control. Immunoprecipitation was confirmed by western blotting. The immunoprecipitates were washed three times with TBS and then with Ser/Thr assay buffer (Ser/Thr Phosphatase Assay Kit 1). We utilized 1 mM phosphoserine peptide generated by (Biomatik, (Canada)) with the sequence, FRTP(pSer)LPQR, corresponding to the S96 region of Miranda. The phosphatase assay was carried out using the Ser/Thr Phosphatase Assay Kit 1 (Millipore Sigma (MA, USA)) following the manufacturer’s protocol, with the incubation time of the phosphatase with the peptide increased to 2 hours.

### Immunoblotting

Lysates/Immunoprecipitates were mixed with SDS sample buffer, separated by SDS-PAGE, transferred to an Immersion-P^SQ^ PVDF membrane (Millipore Sigma) and blotted with the indicated antibodies. Antibodies were HA (Rb; Invitrogen; 1:1000), PP4-19c (Rb; 1:1000), Flfl (Rt; 1:1000), GFP (Rb; Lifetech; 1:1000), HRP-Rt (Jackson Immunoresearch; 1:2000), and HRP-Rb (Cell Signaling Technology; 1:2000). Blots were visualized with SuperSignal West Pico or Femto PLUS Chemoluminescent Substrate (ThermoFisher Scientific) on a BioRad Chemidoc MP Imaging System.

### PCR and quantitative real-time PCR

For PCRs, 3^rd^ instar larval brains were dissected, and RNA was purified using the Qiagen RNeasy kit (Qiagen, Germany). cDNA libraries were generated using M-MLV RT [H-] Point mutant (Promega, WI, USA) according to the manufacturer’s directions. PCR was carried out using DreamTaq 2x Master Mix (Thermo Scientific). Primers were generated by IDT (IA, USA). Products were run on a 1.5% agarose (Fisher) gel in TAE buffer. Primer sequences were: eEF1a1 F (5’-ATGGGCAAGGAAAAGATTCAC-3’); eEF1a1 R (5’-CTACTTCTTGCCCTTGGTGGC-3’); FlflI9 F (5’-ACTTCGTCATCCTCGTCTCTG-3’); FlflI9 R (5’-AGTACTCATCCTCCTCGTAGT-3’); FlflI2 F (5’-CCTATGTGGAAAGACTAAAAG-3’); FlflI2 R (5’-AACTCAGGGCCAGATCAAAG-3’); FlflI4 F (5’-CGTAGAGTTCTCGCCTCTGG-3’); and FlflI4 R (5’-CCTTCTCGGTGAGCATGTTT-3’).

RNA for quantitative Real-time PCR was extracted from 3^rd^ instar larval brains using the Qiagen RNeasy kit, digested with DNase I (NEB) following the manufacturer’s protocol, and repurified with the Qiagen RNeasy kit. Primers were designed using Primer3web version 4.1.0 (http://bioinfo.ut.ee/primer3) as recommended by the manufacturer and generated by IDT. Primer sequences were: eEF1a1 F (5’-CTACAAGTGCGGTGGTATCG-3’); eEF1a1 R (5’-TTATCCAAAACCCAGGCGTA-3’); Flfl Spliced F (5’-CTCCAGCTGTGTCCAGTCC-3’); Flfl Unspliced F (5’-TGTGACGATTAAAGTGCACGA-3’); and Flfl R (5’-TCGTAGTCATCTTCGCCAGA-3’). Optimal concentrations of each primer and primer efficiency were determined, and products were run on an agarose gel to ensure correct products were generated. Real-time PCR reactions were set up using the iTaq Universal Sybr Green One-Step RT-PCR kit (Biorad) according to manufacturer’s directions. Reactions were run in triplicate with a BioRad CFX 384 Real Time PCR system. Results were analyzed using the Delta-Delta Ct and RIV (Camacho Londoño & Philipp, 2016) methods. The expression of each transcript was normalized to eEF1a1 (Ponton *et al*, 2011).

### RNA Immunoprecipitation

Embryos expressing Kin17-HA under the control of *insc-Gal4* were collected for 16 hours, dechorionated with 50% bleach, homogenized with a pestle, and lysed in lysis buffer (150 mM KCl, 20 mM Hepes, pH 7.4, 1mM MgCl2, 0.1% Triton, HALT Protease Inhibitor Cocktail (ThermoFisher Scientific), 40 units/ml Ribolock RNase Inhibitor (ThermoFisher Scientific)). One milligram of protein was incubated with 4 µg of antibody (IgG (Rb, Invitrogen) or HA (Rb, Invitrogen)) and 50 ul of Protein A/G Plus agarose beads (Santa Cruz Biotechnology) for 4 hours. Prior to immunoprecipitation, the beads were pre-blocked overnight with single-stranded Salmon Sperm DNA (0.5 mg/ml, Sigma Aldrich). Beads were collected, washed 3 times with wash buffer, resuspended in NT2 buffer (50 mM Tris (pH 7.4), 150 mM NaCl, 1 mM MgCl2) supplemented with 80 U RNase Inhibitor and 30 ug Proteinase K (Sigma-Aldrich (St. Louis, MO)), and incubated at 55°C for 30 minutes. RNA was purified using RNAzol RT (Molecular Research Center, Inc. (OH, USA)) following the manufacturer’s protocol. RNA was then treated with DNase I (NEB) following the manufacturer’s protocols and 250-500 ng of RNA was used to generate a cDNA library with MMLV H-Reverse transcriptase (Promega) and Oligo dT15 primers (Promega) following the manufacturer’s protocol. PCRs were performed using DreamTaq Hot Start Green PCR Master Mix (ThermoFisher Scientific) in 10 ul volume with 1 ul of the RT reaction with the following primers: Flfl Fwd 1 (5’-CCTATGTGGAAAGACTAAAAG-3’); Flfl Rev 1 (5’-AACTCAGGGCCAGATCAAAG-3’); Flfl Fwd 2 (5’-CGTAGAGTTCTCGCCTCTGG-3’); Flfl Rev 2 (5’-CCTTCTCGGTGAGCATGTTT-3’); Flfl Fwd 3 (5’-CAACGGCCGATACAACCTTCT-3’); Flfl Rev 3 (5’-ATCTTGTCGCGGTCCTTAAGC-3’); Flfl Fwd 4 (5’-ACTTCGTCATCCTCGTCTCTG-3’); and Flfl Rev 4 (5’-CTTTTTGGCCGTCGTCTTGTC-3’). PCR conditions were as follows: 95°C for 5 min, followed by 30 cycles of 95°C for 30 sec, 54°C for 30 sec, 72°C for 30 sec, and a final extension at 72°C for 5 min. Products were detected on a 1.5% agarose gel by staining with SybrSafe (Biorad).

### Quantification and Statistical methods

To quantify the mitotic rate, the ratio of Dpn+, PH3+ neuroblasts to Dpn+ neuroblasts was determined for each brain, with a total of at least 30 neuroblasts counted per brain lobe. The average and standard deviation were calculated from the percentages calculated for each brain.

Quantification of nuclear Pros was determined by colocalization of Pros with Deadpan in interphase cells. Neuroblasts were determined to be positive for nuclear Pros if the nuclear Pros signal was above the cytoplasmic Pros signal. The percentage for each brain was determined by counting at least 25 neuroblasts per brain, and the average and standard deviation were calculated from the percentages calculated for each brain.

To quantify the Flfl levels in each neuroblast, the average intensity of the Flfl signal for 10 neuroblasts from each brain was quantified and the average intensity for each brain was determined. For each experiment, an average was calculated from the average intensity of each brain and the intensity for the mutants was normalized to that of the control.

Statistical analysis was performed using Microsoft Excel. Pairwise comparisons were made using two-tailed, paired or unpaired student’s t-tests. Multiple comparisons were performed using two-tailed, paired or unpaired t-tests with the Bonferroni correction.

## Acknowledgments

We thank Drs. J. Januschke, Z. Lipinszki, C. Doe, and Y.N. Jan for antibodies and fly lines; the Bloomington *Drosophila* Stock Center and the TRiP at Harvard Medical School (NIH/NIGMS R01-GM084947) for providing transgenic RNAi fly stocks used in this study; Zhu Lab members for thoughtful discussion and comments; Best Gene, Inc for generating transgenic flies.

This work was supported by the National Institute of Neurological Disorders and Stroke of the National Institutes of Health under Award Number R01NS085232 (S.Z.) and R21NS109748 (S.Z.). The authors declare no competing financial interests.

## Author Contributions

Conceptualization and experimental design were performed by M. Connell, Y. Xie, and S. Zhu. Experiments were performed by M. Connell, Y. Xie, and R. Chen. Funding was obtained by S. Zhu. Experimental data were analyzed by M. Connell. The paper was written by M. Connell and S. Zhu.

## References

Albertson R & Doe CQ (2003) Dlg, Scrib and Lgl regulate neuroblast cell size and mitotic spindle asymmetry. Nat Cell Biol 52: 166–170

Angulo JF, Rouer E, Mazin A, Mattei MG, Tissier A, Horellou P, Benarous R & Devoret R (1991) Identification and expression of the cDNA of KIN17, a zinc-finger gene located on mouse chromosome 2, encoding a new DNA-binding protein. Nucleic Acids Res 192: 5117–5123

Atwood SX, Chabu C, Penkert RR, Doe CQ & Prehoda KE (2007) Cdc42 acts downstream of Bazooka to regulate neuroblast polarity through Par-6 aPKC. J Cell Sci 1202: 3200–3206

Atwood SX & Prehoda KE (2009) Phosphorylation-mediated cortical displacement of fate determinants by aPKC during neuroblast asymmetric cell division. Curr Biol 192: 723–729

Bailey MJ & Prehoda KE (2015) Establishment of Par-polarized cortical domains via phosphoregulated membrane motifs. Dev Cell 352: 199–210

Bello B, Reichert H & Hirth F (2006) The brain tumor gene negatively regulates neural progenitor cell proliferation in the larval central brain of Drosophila. Development 1332: 2639–2648

Betschinger J, Mechtler K & Knoblich JA (2003) The Par complex directs asymmetric cell division by phosphorylating the cytoskeletal protein Lgl. Nature 4222: 326–330

Betschinger J, Mechtler K & Knoblich JA (2006) Asymmetric segregation of the tumor suppressor Brat regulates self-renewal in Drosophila neural stem cells. Cell 1242: 1241–1253

Bier E, Vaessin H, Younger-Shepherd S, Jan LY & Jan YN (1992) deadpan, an essential pan-neural gene in Drosophila, encodes a helix-loop-helix protein similar to the hairy gene product. Genes Dev 62: 2137–2151

Brand AH & Perrimon N (1993) Targeted gene expression as a means of altering cell fates and generating dominant phenotypes. Development 1182: 401–415

Camacho Londoño J & Philipp SE (2016) A reliable method for quantification of splice variants using RT-qPCR. BMC Mol Biol 172: 1–12

Colonques J, Ceron J, Reichert H & Tejedor FJ (2011) A transient expression of Prospero promotes cell cycle exit of Drosophila postembryonic neurons through the regulation of Dacapo. PLoS One 62: 1–13

Doe CQ, Chu-LaGraff Q, Wright DM & Scott MP (1991) The prospero gene specifies cell fates in the Drosophila central nervous system. Cell 652: 451–464

Gaspar VP, Ramos AC, Cloutier P, Pattaro Junior JR, Duarte Junior FF, Bouchard A, Seixas FAV, Coulombe B & Fernandez MA (2021) Interactome analysis of KIN (Kin17) shows new functions of this protein. Curr Issues Mol Biol 432: 767–781

Gómez-López S, Lerner RG & Petritsch C (2014) Asymmetric cell division of stem and progenitor cells during homeostasis and cancer. Cell Mol Life Sci 712: 575–597

Hannaford MR, Ramat A, Loyer N & Januschke J (2018) aPKC-mediated displacement and actomyosin-mediated retention polarize Miranda in Drosophila neuroblasts. Elife 72: 1–22

Helps NR, Brewis ND, Lineruth K, Davis T, Kaiser K & Cohen PTW (1998) Protein Phosphatase 4 is an essential enzyme required for organisation of microtubules at centrosomes in Drosophila embryos. J Cell Sci 1112: 1331–1340

Herold N, Will CL, Wolf E, Kastner B, Urlaub H, Lu R & Luhrmann R (2009) Conservation of the protein composition and electron microscopy structure of Drosophila melanogaster and human spliceosomal complexes. Mol Cell Biol 292: 281–301

Hirata J, Nakagoshi H, Nabeshima YI & Matsuzaki F (1995) Asymmetric segregation of the homeodomain protein Prospero during Drosophila development. Nature 3772: 627–630

Homem CCF & Knoblich JA (2012) Drosophila neuroblasts: a model for stem cell biology. Development 1392: 4297–4310

Ikeshima-Kataoka H, Skeath JB, Nabeshima YI, Doe CQ & Matsuzaki F (1997) Miranda directs Prospero to a daughter cell during Drosophila asymmetric divisions. Nature 3902: 625–629

Irion U, Adams J, Chang CW & St Johnston D (2006) Miranda couples oskar mRNA/Staufen complexes to the bicoid mRNA localization pathway. Dev Biol 2972: 522–533

Knoblich JA (2010) Asymmetric cell division: recent developments and their implications for tumor biology. Nat Rev Mol Cell Biol 112: 849–860

Knoblich JA, Jan LY & Jan YN (1995) Asymmetric segregation of Numb and Prospero during cell division. Nature 3772: 624–627

Kou WZ, Xu SL, Wang Y, Wang LW, Wang L, Chai XY & Hua QL (2014) Expression of Kin17 promotes the proliferation of hepatocellular carcinoma cells in vitro and in vivo. Oncol Lett 82: 1190–1194

Lai SL & Doe CQ (2014) Transient nuclear Prospero induces neural progenitor quiescence. Elife 32: 1–12

Lee CY, Wilkinson BD, Siegrist SE, Wharton RP & Doe CQ (2006) Brat is a Miranda cargo protein that promotes neuronal differentiation and inhibits neuroblast self-renewal. Dev Cell 102: 441–449

Lipinszki Z, Lefevre S, Savoian MS, Singleton MR, Glover DM & Przewloka MR (2015) Centromeric binding and activity of Protein Phosphatase 4. Nat Commun 62: 1–13

Masson C, Menaa F, Pinon-Lataillade G, Frobert Y, Chevillard S, Radicella JP, Sarasin A & Angulo JF (2003) Global genome repair is required to activate KIN17, a UVC-responsive gene involved in DNA replication. Proc Natl Acad Sci U S A 1002: 616–621

Matsuzaki F, Ohshiro T, Ikeshima-Kataoka H & Izumi H (1998) Miranda localizes Staufen and Prospero asymmetrically in mitotic neuroblasts and epithelial cells in early Drosophila embryogenesis. Development 1252: 4089–98

Mendoza MC, Booth EO, Shaulsky G & Firtel RA (2007) MEK1 and Protein Phosphatase 4 Coordinate Dictyostelium Development and Chemotaxis. Mol Cell Biol 272: 3817–3827

Miccoli L, Frouin I, Novac O, Paola DD, Haroer F, Zannis-Hadjopoulos M, Maga G, Biard DSF & Angulo JF (2005) The human stress-activated protein Kin17 belongs to the multiprotein DNA replication complex and associates in vivo with mammalian replication origins. Mol Cell Biol 252: 3814–3830

Mollinari C, Lange B & González C (2002) Miranda, a protein involved in neuroblast asymmetric division, is associated with embryonic centrosomes of Drosophila melanogaster. Biol Cell 942: 1–13

Ni JQ, Liu LP, Binari R, Hardy R, Shim HS, Cavallaro A, Booker M, Pfeiffer BD, Markstein M, Wang H, et al (2009) A drosophila resource of transgenic RNAi lines for neurogenetics. Genetics 1822: 1089–1100

Otake LR, Scamborova P, Hashimoto C & Steitz JA (2002) The divergent U12-type spliceosome is required for pre-mRNA splicing and is essential for development in Drosophila. Mol Cell 92: 439–446

Ou Y & Rattner J (2004) The centrosome in higher organisms: structure, composition, and duplication. Int Reivew Cytyology 2382: 119–182

Petritsch C, Tavosanis G, Turck CW, Jan LY & Jan YN (2003) The Drosophila myosin VI Jaguar is required for basal protein targeting and correct spindle orientation in mitotic neuroblasts. Dev Cell 42: 273–281

Petronczki M & Knoblich JA (2001) DmPAR-6 directs epithelial polarity and asymetric cell division of neuroblasts in Drosophila. Nat Cell Biol 32: 43–49

Pinon-Lataillade G (2004) KIN17 encodes an RNA-binding protein and is expressed during mouse spermatogenesis. J Cell Sci 1172: 3691–3702

Ponton F, Chapuis MP, Pernice M, Sword GA & Simpson SJ (2011) Evaluation of potential reference genes for reverse transcription-qPCR studies of physiological responses in Drosophila melanogaster. J Insect Physiol 572: 840–50

Rappsilber J, Ryder U, Lamond AI & Mann M (2002) Large-scale proteomic analysis of the human spliceosome. Genome Res 132: 1231–1245

Rothwell WF & Sullivan W. (2000) Fluorescent Analysis of Drosophila Embryos. Drosoph Protoc: 141–157 [PREPRINT]

Ruan L, Jiang W & Zhang H (2018) Relationships of Kin17 protein expression with clinical features and prognosis of colorectal cancer. Transl Cancer Res 72: 1072–1078

Schindelin J, Arganda-Carrera I, Frise E, Verena K, Mark L, Tobias P, Stephan P, Curtis R, Stephan S, Benjamin S, et al (2009) Fiji - an Open Source platform for biological image analysis. Nat Methods 9

Schober M, Schaefer M & Knoblich JA (1999) Bazooka recruits inscuteable to orient asymmetric cell divisions in Drosophila neuroblasts. Nature 4022: 548–551

Schuldt AJ, Adams JHJ, Davidson CM, Micklem DR, Haseloff J, St. Johnston D & Brand AH (1998) Miranda mediates asymmetric protein and RNA localization in the developing nervous system. Genes Dev 122: 1847–1857

Shen CP, Jan LY & Jan YN (1997) Miranda is required for the asymmetric localization of prospero during mitosis in Drosophila. Cell 902: 449–458

Siller KH, Cabernard C & Doe CQ (2006) The NuMA-related Mud protein binds Pins and regulates spindle orientation in Drosophila neuroblasts. Nat Cell Biol 82: 594–600

Song Y & Lu B (2012) Interaction of notch signaling modulator numb with α-adaptin regulates endocytosis of notch pathway components and cell fate determination of neural stem cells. J Biol Chem 2872: 17716–17728

Sousa-Nunes R, Chia W & Somers WG (2009) Protein Phosphatase 4 mediates localization of the Miranda complex during Drosophila neuroblast asymmetric divisions. Genes Dev 232: 359–372

Spradling AC, Stern D, Beaton A, Rhem EJ, Laverty T, Mozden N, Misra S & Rubin GM (1999) The Berkeley Drosophila Genome Project gene disruption project: Single P-element insertions mutating 25% of vital Drosophila genes. Genetics 1532: 135–177

Wodarz A, Ramrath A, Kuchinke U & Knust E (1999) Bazooka provides an apical cue for inscuteable localization in Drosophila neuroblasts. Nature 4022: 544–547

Zeng T, Gao H, Yu P, He H, Ouyang X, Deng L & Zhang Y (2011) Up-regulation of Kin17 is essential for proliferation of breast cancer. PLoS One 62: 1–10

Zhang F, Huang Z-X, Bao H, Cong F, Wang H, Chai PC, Xi Y, Ge W, Somers WG, Yang Y, et al (2016) Phosphotyrosyl phosphatase activator facilitates localization of Miranda through dephosphorylation in dividing neuroblasts. Development 1432: 35–44

Zhang Y, Gao H, Gao X, Huang S, Wu K, Yu X, Yuan K & Zeng T (2017a) Elevated expression of Kin17 in cervical cancer and its association with cancer cell proliferation and invasion. Int J Gynecol Cancer 272: 628–633

Zhang Y, Huang S, Gao H, Wu K, Ouyang X, Zhu Z, Yu X & Zeng T (2017b) Upregulation of KIN17 is associated with non-small cell lung cancer invasiveness. Oncol Lett 132: 2274–2280

