## Supplementary Figures 1-8 for "Kin17 drives dissociation of Mira from the centrosome in neuroblasts by regulating splicing of Flfl"

**Supplementary Figure legends**

**Supplementary Figure 1. Mira Cortical localization is affected by Kin17 knockdown.** (**A**) Quantification of cortical/cytoplasmic localization of Mira in in wild type, *pros*$^{17/+}$, Kin17 RNAi *pros* neuroblasts. Interphase, n=74 NBs (ctrl), 101 (*pros*$^{17/+}$), 101 (*Kin17 RNAi; pros*$^{17/+}$); Prophase, n=63 (ctrl), 57 (*pros*$^{17/+}$), 85 (*Kin17 RNAi; pros*$^{17/+}$); Prometaphase/Metaphase, n=57 (ctrl), 56 (*pros*$^{17/+}$), 98 (*Kin17 RNAi; pros*$^{17/+}$); Anaphase/Telophase, n=35 (ctrl), 18 (*pros*$^{17/+}$), 26 (*Kin17 RNAi*; *pros*$^{17/+}$). (**B**) Cytoplasmic and cortical localization of Mira in *Kin17 RNAi; pros*$^{17/+}$ neuroblasts. Metaphase depicts normal (left) and weak (right) basal cortical localization of Mira. Scale bars, 5 µm. (**C**) Quantification of apical and/or basal centrosomal localization of Mira in neuroblasts expressing *Kin17 RNAi* . Prophase, n=37 NBs (ctrl), 30 (*pros*$^{17/+}$), 50 (*Kin17 RNAi; pros17/+*); Metaphase, n=17 (ctrl), 12 (*pros*$^{17/+}$), 56 (*Kin17 RNAi; pros*$^{17/+}$). (**D**) Mira localizes to the centrosome in neuroblasts expressing *Kin17 RNAi* at interphase. Arrows indicate centrosomes. Scale bars, 5 µm. (**E**) Quantification of Mira localization to the centrosome in neuroblasts expressing *Kin17 RNAi* (n=65 NBs).

**Supplementary Figure 2. Kin17 knockdown does not affect Numb localization.** (**A**) Localization of Numb in control and *Kin17 RNAi* neuroblasts at the interphase and metaphase. Arrows indicate centrosomes. Scale bars, 5 µm. (**B**) Quantification of Numb localization during the cell cycle in control and *Kin17 RNAi* neuroblasts. Interphase, n=107 NBs (control), 87 (*Kin17 RNAi*); Pro/Metaphase, n=105 (control), 35 (*Kin17 RNAi*).

**Supplementary Figure 3. Phosphorylation status at S96 affect Mira localization in neuroblasts.** (**A**) Quantification of Mira localization to the cortex/cytoplasm in neuroblasts expressing Mira$^{WT}$, Mira$^{S96D}$, and Mira$^{S96A}$. Interphase, n=119 NBs (Mira$^{WT}$), 151 (Mira$^{S96D}$), 135 (MiraS96A); Prophase, n=50 (Mira$^{WT}$), 30 (Mira$^{S96D}$), 52 (Mira$^{S96A}$); Prometaphase/Metaphase, n=70 (Mira$^{WT}$), 52 (Mira$^{S96D}$), 94 (MiraS$^{S96A}$); Anaphase/Telophase, n=41 (Mira$^{WT}$), 30 (Mira$^{S96D}$), 67 (Mira$^{S96A}$). (**B**) Quantification of Mira localization to the apical and/or basal centrosomes in neuroblasts expressing Mira$^{WT}$, Mira$^{S96D}$, and Mira$^{S96A}$. Prophase, n=34 NBs (Mira$^{WT}$), 58 (Mira$^{S96D}$), 9 (Mira$^{S96A}$); Metaphase, n=9 (Mira$^{WT}$), 37 (Mira$^{S96D}$), 2 (Mira$^{S96A}$).

**Supplementary Figure 4. Phosphorylation at T591 does not lead to centrosome localization of Mira.** (**A**) Centrosomal localization of Mira$^{WT}$, Mira$^{T591D}$, and Mira$^{T591A}$ constructs. Scale bar, 5 µm. (**B**) Quantification of centrosomal localization of Mira$^{WT}$, Mira$^{T59D}$, and Mira$^{T591A}$ constructs.

Interphase, n=51 NBs (Mira$^{WT}$), 70 (Mira$^{T591D}$), 64 (Mira$^{T591A}$); Prophase, n=41 (Mira$^{WT}$), 71 (Mira$^{T591D}$), 32 (Mira$^{T591A}$); Prometaphase/Metaphase, n=44 (Mira$^{WT}$), 46 (Mira$^{T591D}$), 57 (Mira$^{T591A}$); Anaphase/Telophase, n=28 (Mira$^{WT}$), 28 (Mira$^{T591D}$), 18 (Mira$^{T591A}$).

**Supplementary Figure 5. Reduction of aPKC does not rescue Kin17 Phenotype.** (**A**) Bazooka localization in control and *Kin17 RNAi* neuroblasts (outlined by dotted circles). Scale bars, 5 µm. (**B**) aPKC localization in control and *Kin17 RNAi* neuroblasts. Scale bars, 10 µm. (**C**) Kin17 knockdown still leads to nuclear localization of Pros in *aPKC*$^{K06403/+}$ heterozygous mutant larval brains. Arrows indicate nuclear Pros. Scale bars, 50 µm. (**D**) Quantification of nuclear Pros localization in *Kin17 RNAi* (n=7 brains) and *Kin17 RNAi*, *aPKC*$^{K06403/+}$ (n=8) larval neuroblasts. Error bars, 1 SD; n.s., not significant.

**Supplementary Figure 6. PP4 regulates Mira localization likely by dephosphorylating Mira at S96.** (**A**) Quantification of cortical/cytoplasmic localization of Mira in *PP4-19c* and *Flfl RNAi* neuroblasts. Interphase, n=200 NBs (*PP4-19c RNAi*), 136 (*Flfl RNAi*); Prophase, n=32 (*PP4-19c RNAi*), 98 (*Flfl RNAi*); Prometaphase/Metaphase, n=81 (*PP4-19c RNAi*), 132 (*Flfl RNAi*); Anaphase/Telophase, n=32 (*PP4-19c RNAi*), 32 (*Flfl RNAi*). (**B**) Quantification of apical and/or basal centrosomal localization of Mira in *PP4-19c* and *Flfl RNAi* neuroblasts. Prophase, n=25 NBs (*PP4-19c RNAi*), 73 (*Flfl RNAi*); Metaphase, n=51 (*PP4-19c RNAi*),83 (*Flfl RNAi*). (**C**) Nuclear localization of Pros in control (n=4 brains) and *flfl*$^{n42}$ brains. Dotted lines indicate neuroblasts. Scale bar, 5 µ m. (**D**) Quantification of Pros localization in control (n=4 brains) and *flfl*$^{n42}$ (n=8) mutant neuroblasts. (**E**) PH3 staining in control and *flfl*$^{n42}$ mutant neuroblasts. Arrows indicate mitotic cells. Scale bar, 25 µm. (**F**) Quantification of mitotic rate in control (n=10) and *flfl*$^{n42}$ (n=23) brains. Error bars, 1 SD. ***, $p<0.001$. (**G**) Cytoplasmic and cortical localization of Mira$^{S96D}$ in the *Flfl RNAi* background. Metaphase depicts normal (left) and weak (right) basal cortical localization. Scale bars, 5 µm. (**H**) Quantification of cortical/cytoplasmic localization of Mira$^{WT}$ and Mira$^{S96D}$ in the *Flfl RNAi* background. Interphase, n=40 NBs (Mira$^{WT}$), 52 (Mira$^{S96D}$), 130 NBs (Mira$^{WT}$, *Flfl RNAi*), 117 (Mira$^{S96D}$, *Flfl RNAi*); Prophase, n=25 (Mira$^{WT}$), 28 (Mira$^{S96D}$), 69 (Mira$^{WT}$, *Flfl RNAi*), 51 (Mira$^{S96D}$, *Flfl RNAi*); Prometaphase/Metaphase, n=26 (Mira$^{WT}$), 37 (Mira$^{S96D}$), 65 (Mira$^{WT}$, *Flfl RNAi*), 53 (Mira$^{S96D}$, *Flfl RNAi*); Anaphase/Telophase, n=21 (Mira$^{WT}$), 31 (Mira$^{S96D}$), 42 (Mira$^{WT}$, *Flfl RNAi*), 32 (Mira$^{S96D}$, *Flfl RNAi*).

**Supplementary Figure 7. Mira localization in neuroblasts expressing RFP-Flfl in the Kin17 RNAi background** (**A**) Cytoplasmic and cortical localization of Mira in *Kin17 RNAi*; RFP-Flfl neuroblasts. Metaphase depicts normal (left) and weak (right) basal cortical localization. Scale bars, 5 µm. (**B**) Quantification of Mira localization to the cortex/cytoplasm in neuroblasts expressing RFP-Flfl in the *Kin17 RNAi* background throughout the cell cycle. Interphase, n=102 NBs (RFP-Flfl), 46 (*Kin17 RNAi*; RFP-Flfl), 49 (*Kin17 RNAi*); Prophase, n=62 (RFP-Flfl), 34 (*Kin17 RNAi,* RFP-Flfl), 25 (*Kin17 RNAi*); Prometaphase/Metaphase, n=65 (RFP-Flfl), 37 (*Kin17 RNAi,* RFP-Flfl), 27 (*Kin17 RNAi*); Anaphase/Telophase, n=31 (RFP-Flfl), 17 (*Kin17 RNAi*, RFP-Flfl), 11 (*Kin17 RNAi*). (**C**) Quantification of Mira to the centrosome in neuroblasts expressing RFP-Flfl in the *Kin17 RNAi* background throughout the cell cycle. Prophase, n=33 NBs (RFP-Flfl), 23 (*Kin17 RNAi*, RFP-Flfl), 16 (*Kin17 RNAi*); Metaphase, n=21 NBs (RFP-Flfl), 16 (*Kin17 RNAi*, RFP-Flfl), 15 (*Kin17 RNAi*). (**D**) RFP-Flfl rescues PP4-19c localization in *Kin17 RNAi* neuroblasts. Scale bars, 5 µm. (**E**) Quantification of nuclear PP4-19c localization. n= 7 brains (Ctrl), 8 ($flfl^{n42}$), 5 (*Kin17 RNAi*), 5 (RFP Flfl), 10 (*Kin17 RNAi,* RFP Flfl).

**Supplementary Figure 8. Mira localization is not affected by the loss of U2A but cortical localization is affected by loss of U6atac.** (**A**) Localization of Mira throughout the cell cycle in U2A RNAi knockdown neuroblasts. Scale bar, 5 µm. (**B**) Quantification of Mira centrosomal localization in U2A RNAi knockdown neuroblasts. n=80 NBs (Interphase); 40 (Prophase); 61 (Prometaphase/Metaphase); 24 (Anaphase/Telophase). (**C**) Quantification of Mira localization to the apical and/or basal centrosome in $u6atac^{k01105}$ ($u6atac^{-/-}$) mutant neuroblasts. Prophase, n=33 NBs; Prometaphase/Metaphase, n=33 NBs. (**D**) Cortical localization of Mira in metaphase $u6atac^{-/-}$ neuroblasts. Arrows indicate basal cortex. Scale bar, 5 µm. (**E**) Quantification of cortical/cytoplasmic localization of Mira throughout the cell cycle in $u6atac^{-/-}$ mutant neuroblasts. n=69 NBs (Interphase), 46 (Prophase), 66 (Prometaphase/Metaphase), 19 (Anaphase/Telophase).

### Connell_Fig. S1

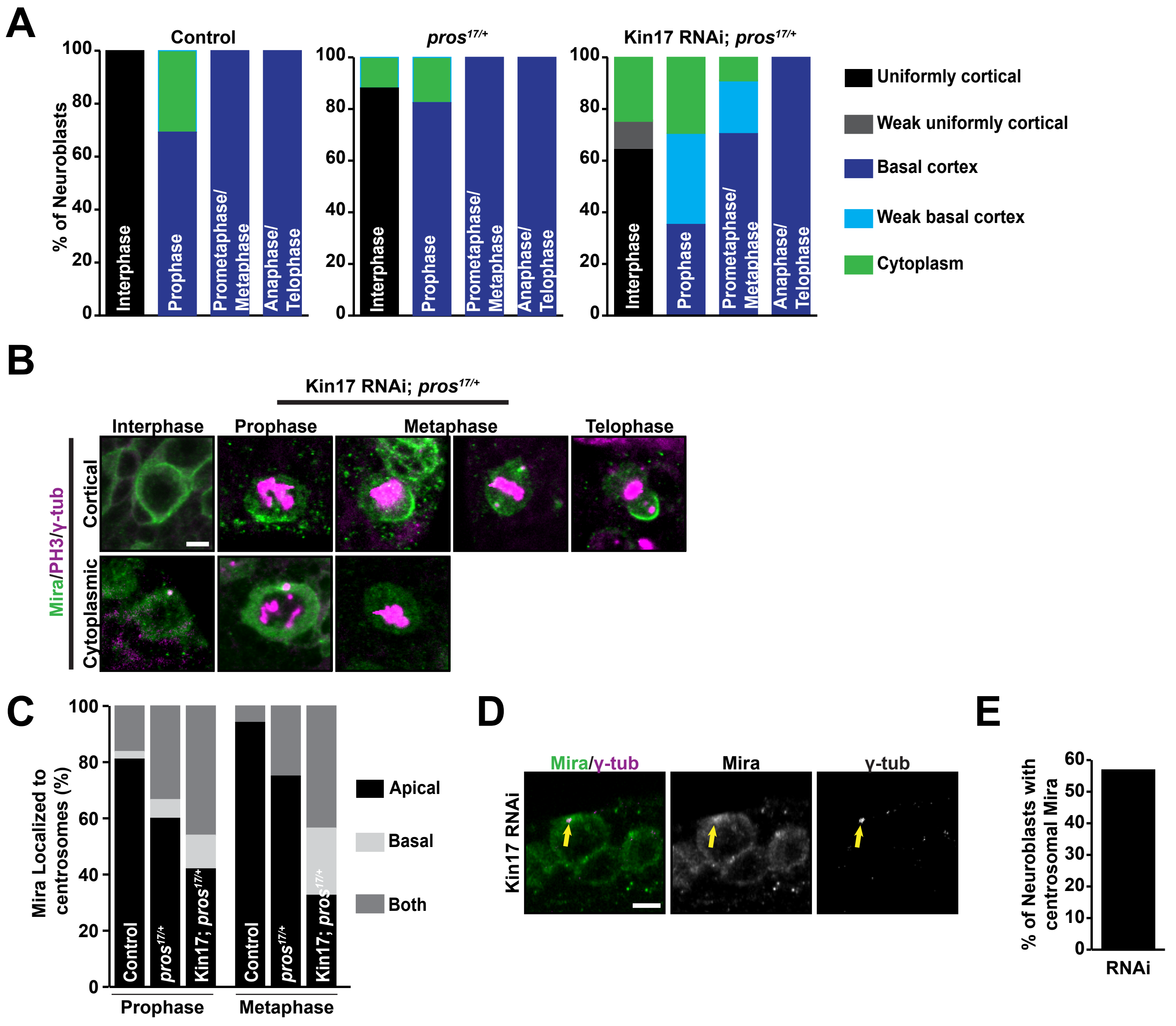

### Connell_Fig. S2

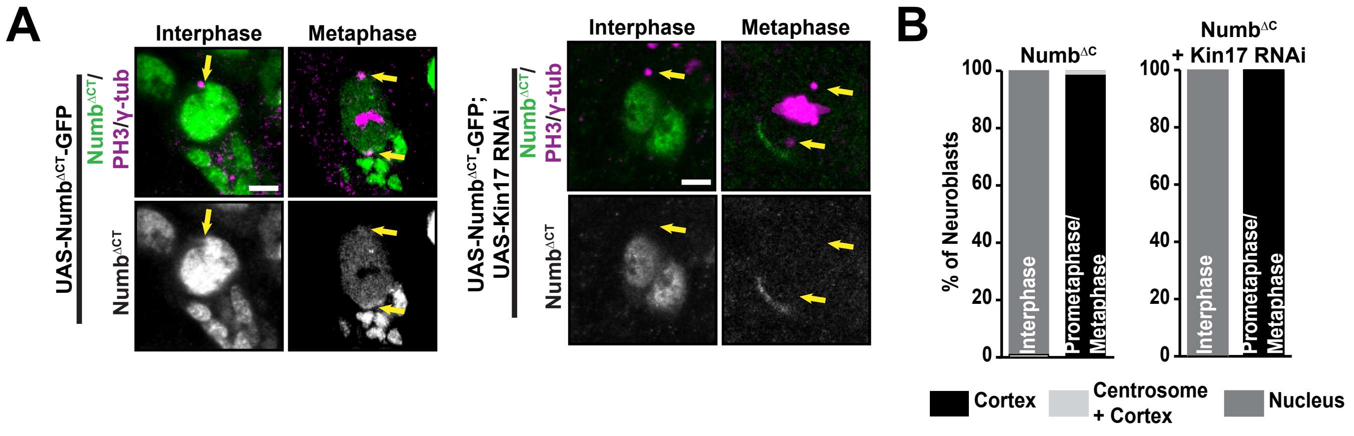

Connell_Fig. S3

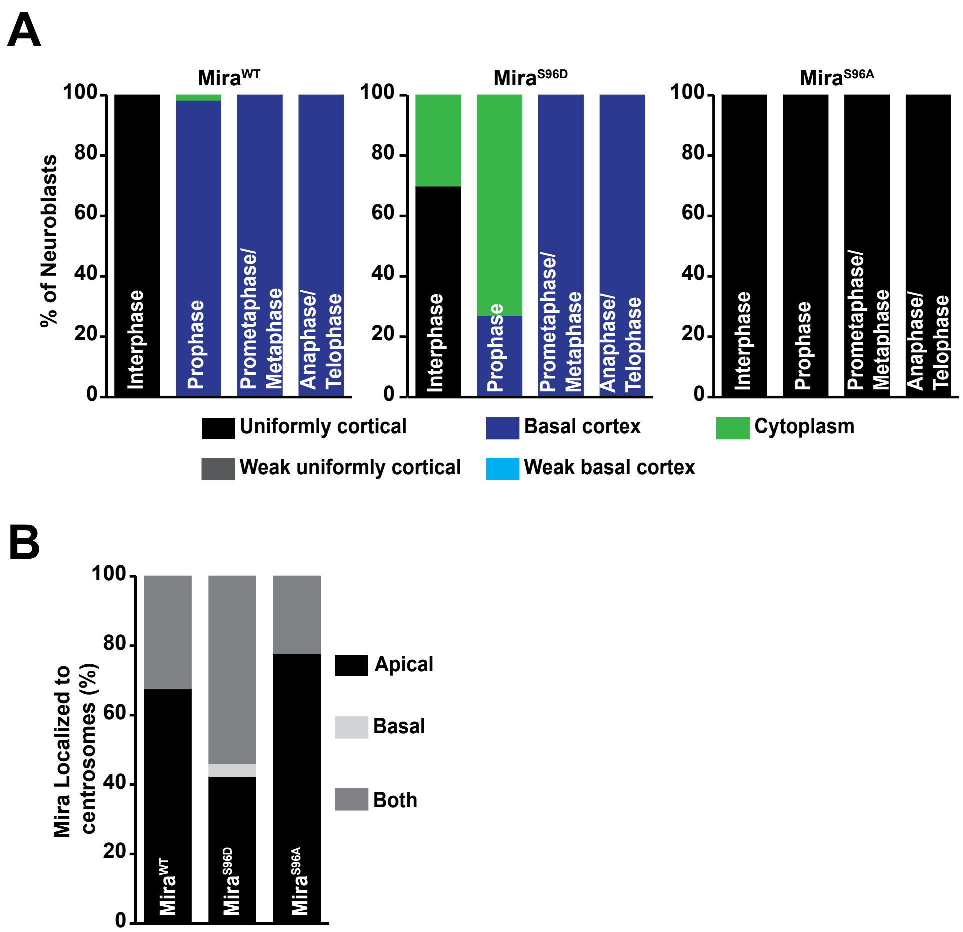

### Connell_Fig. S4

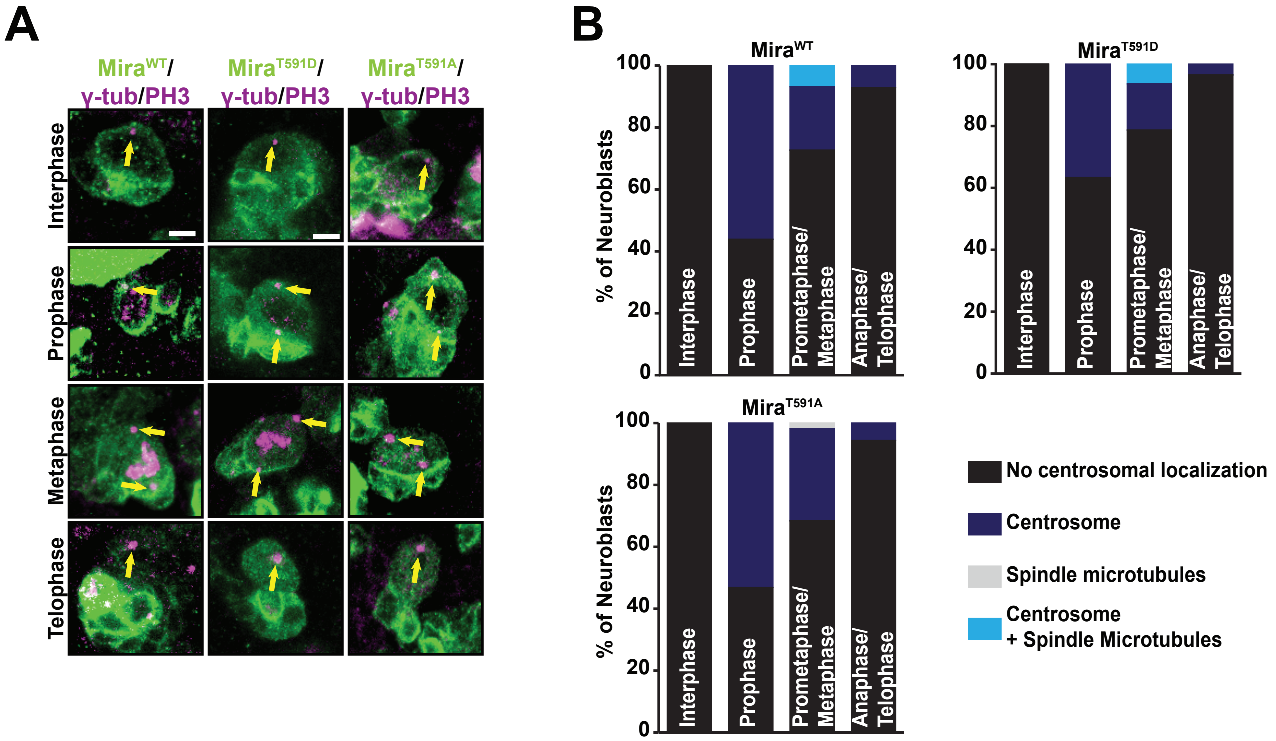

### Connell_Fig. S5

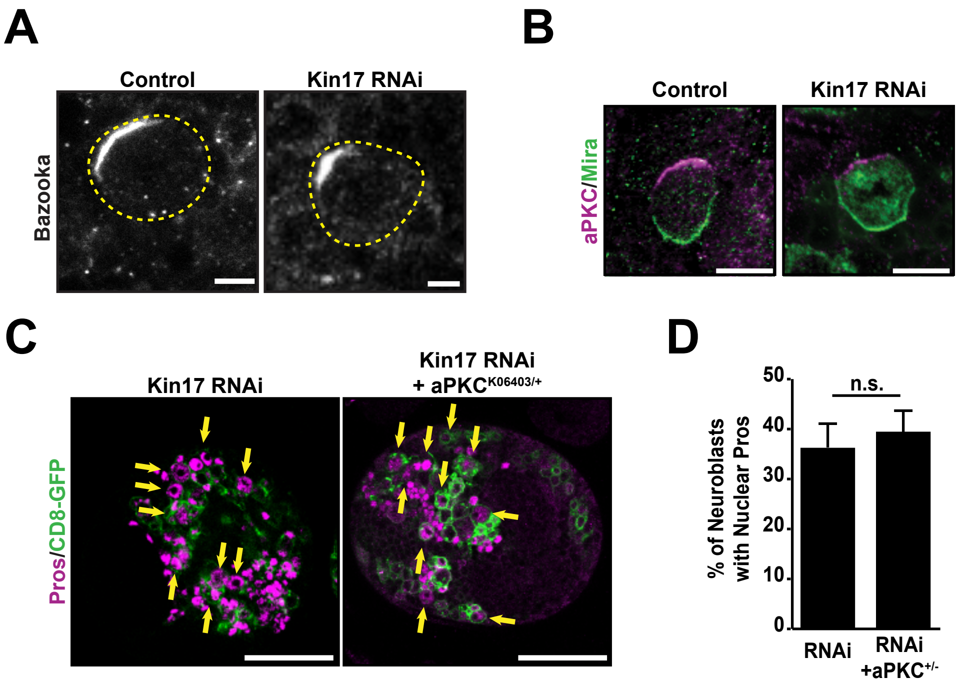

Connell_Fig. S6

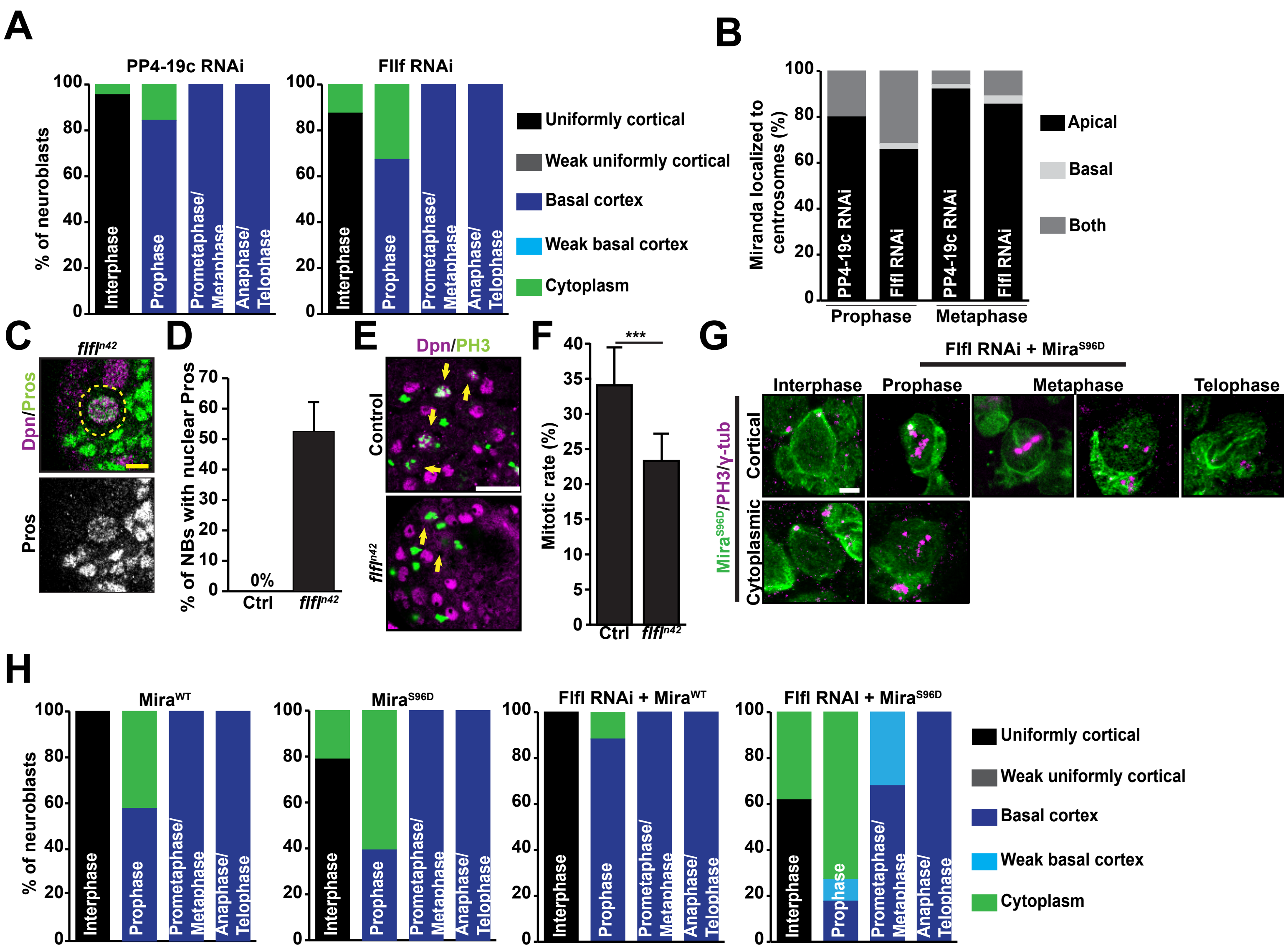

Connell_Fig. S7

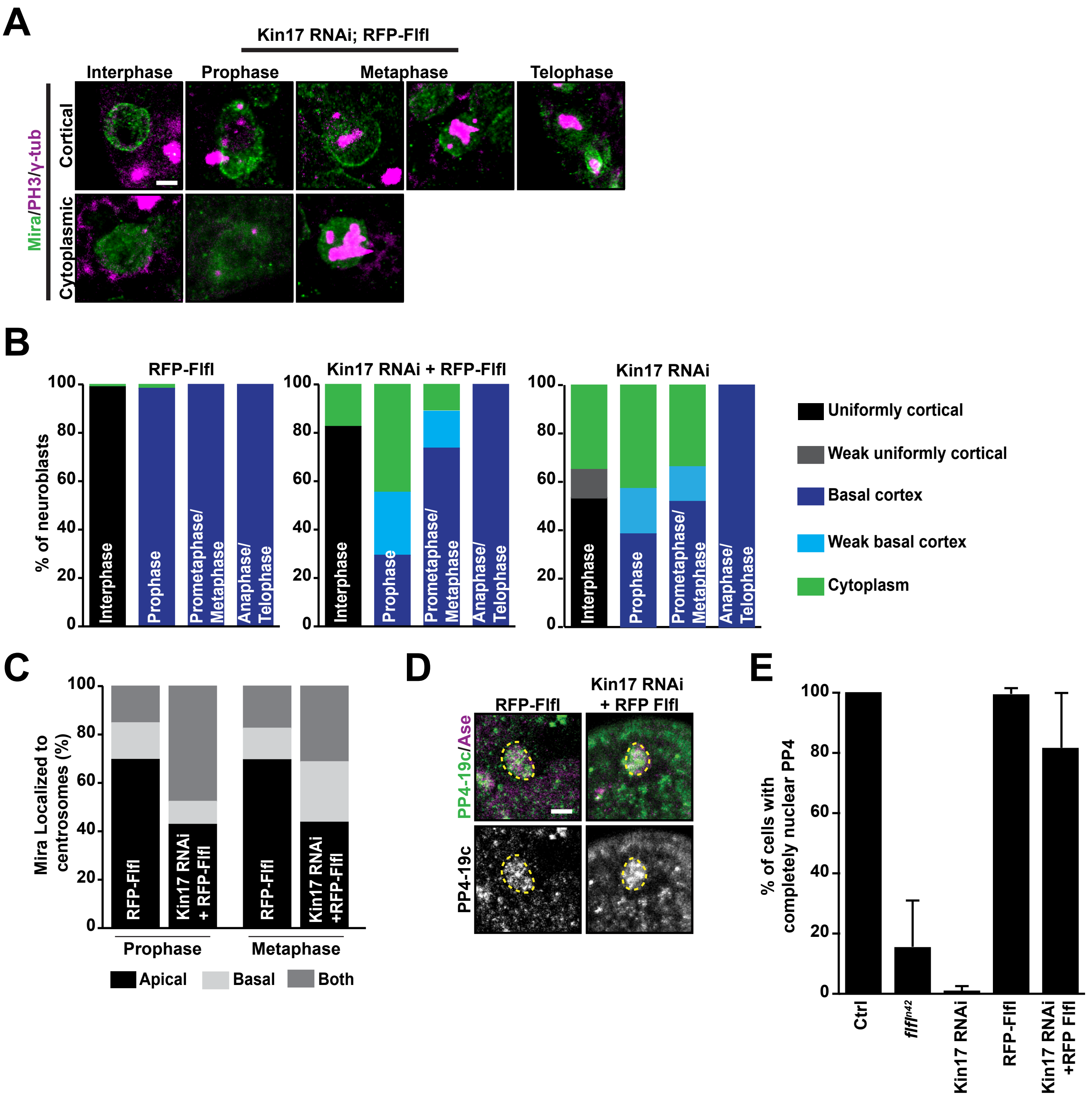

### Connell_Fig. S8

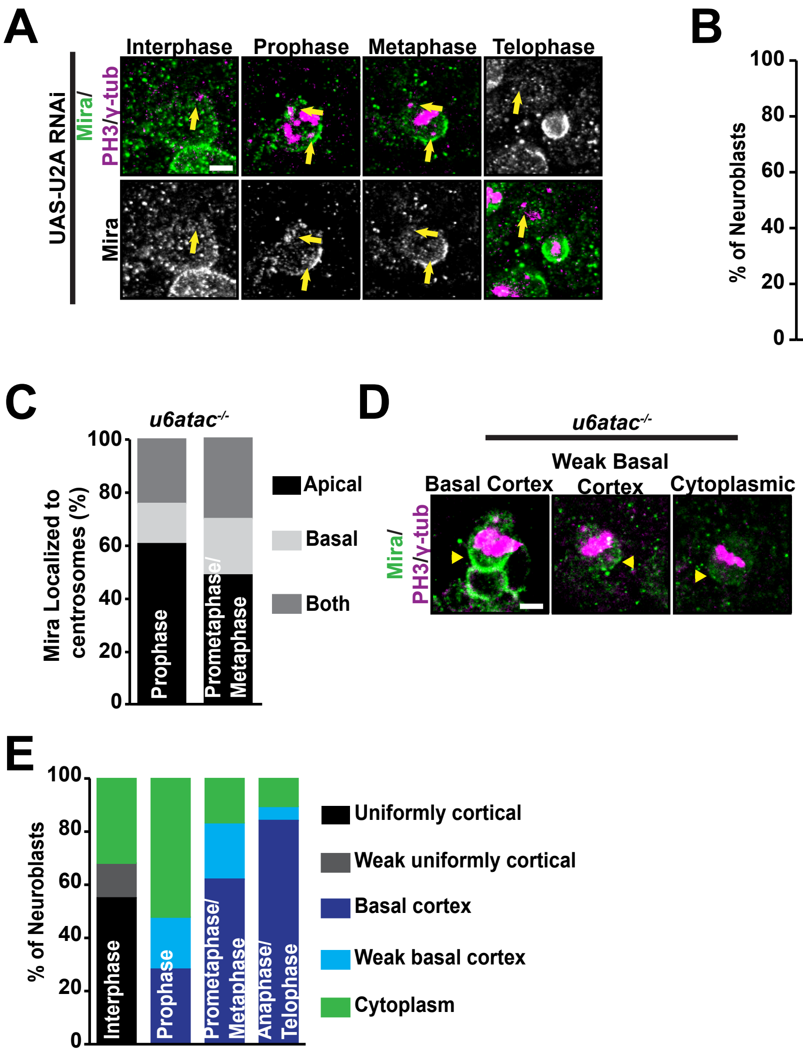
